# Experimental study of modified human tendon stem cells to promote graft ligamentation and tendon-bone healing after anterior cruciate ligament reconstruction

**DOI:** 10.1101/2022.12.14.520376

**Authors:** Haibo Zhao, Jinli Chen, Chao Qi, Tianrui Wang, Tongda Liang, Xiaokun Hao, Xiang Li, Xiangzhi Yin, Tengbo Yu, Yingze Zhang

## Abstract

Restoring the normal structure and function of injured tendons is one of the biggest challenges faced by the Department of Orthopedics and Sports Medicine. Tendon-derived stem cells (TDSCs), a new type of pluripotent stem cells with multidirectional differentiation potential, are expected to be promising cell seeds for the treatment of tendon injury and tendon-bone healing in the future. In this study, tendon stem cells were successfully isolated from human tissues, which were positive for markers CD44, CD90, and CD105, and exhibited clonality and multilineage differentiation ability. Analysis of single-cell sequencing results and mass spectrometry identification results showed that there were differences in protein expression during CTGF-induced TDSC tendon differentiation. Reverse Co-IP, qPCR, WB, and immunofluorescence detection all confirmed that CTGF directly interacts with KIT, thereby mediating the transcription factor HES1 to regulate the Wnt/β-catenin signaling pathway (GSK3β, β-catenin, TCF4). ChIP-qPCR and dual-luciferase reporter gene assays indicated that HES1 regulates stem cell differentiation by directly regulating the expression of GSK3β in the Wnt/β-catenin pathway. Rats were treated with TDSCs overexpressing the KIT gene after repair surgery. This method had a more ideal recovery effect than other methods through animal behavioral scores, mechanical properties testing, and HE staining tissue observation. This study found that the use of modified human tendon stem cells (hTDSC) could promote graft ligamentization and tendon-bone healing after ACL reconstruction, which could provide an effective way for faster and better recovery from tendon injury.

## Introduction

The anterior cruciate ligament (ACL) is a dense band of connective tissue that flows from the femur to the tibia, and its function is to prevent excessive anterior-posterior translation of the knee joint and maintain joint stability [1, 2]. Tear or rupture of the ACL is a very common injury in sports medicine, accounting for more than 50 percent of all knee injuries [3]. ACL reconstruction is the most effective way to treat ACL injury. This way can well restore the stability of the knee joint, to reduce a series of problems such as knee instability caused by ACL rupture, secondary meniscus tear, cartilage injury and knee degenerative changes [4].

The success or failure of ACL reconstruction depends on multiple factors, including the selection of ligament graft, the location of bone canal, the fixation method of graft, the rehabilitation process after reconstruction and so on [5]. Among the many factors, the issue of graft lamentation has attracted much attention. When the tendon graft implanted in the body can achieve “ligamentalization” and become a normal ligament tissue is very important for guiding the formulation of rehabilitation procedures after ACL reconstruction and when it can return to normal exercise level [6, 7].

Tendon-bone healing is a reconstructive procedure that requires tendon grafting to the bone tunnel (eg, ACL reconstruction) or bone surface (eg, rotator cuff tendon repair) following injury to the junction between tendon, ligament, and bone healing [8]. Tendon-bone healing after ACL reconstruction is a complex process that has a significant impact on patient outcomes [9]. After ACL reconstruction, tendon grafts generally go through the process of inflammatory destruction, tissue cell regeneration, graft revascularization, collagen fiber remodeling, and ligament reconstruction, while tendon tissue has fewer microvessels, weaker self-repair ability, and scar tissue. Formation becomes the main way of its healing, resulting in imperfect early postoperative rehabilitation and biomechanical strength of the graft [10, 11]. How to repair tendon injury safely and effectively has become the focus, difficulty and hot issue of early rehabilitation after ACL reconstruction.

Tendon repair using tendon-derived stem cells (TDSCs) is particularly relevant for tendon and tendon-bone junction repair because of their tissue origin [12]. TDSCs were identified within tendon tissue, which possesses universal stem cell characteristics such as clonality, pluripotency, and self-renewal capacity [13]. TDSCs can differentiate into adipocytes, chondrocytes and osteocytes in vitro and form tendon-like, cartilage-like, osteo-like, and tendon-osseous junction-like tissues in rat models [13, 14]. TDSCs are of great significance to tendon tissue development, homeostasis maintenance, injury repair and regeneration, and can reshape the biomechanical function of tendon to a great extent. Therefore, using TDSCs as the cell source of tendon-bone connection repair has more advantages than other mesenchymal stem cells [14]. Therefore, TDSCs may be a potential therapeutic target for tendon-bone healing.

Connective tissue growth factor (CTGF), known as CCN2, is a 38kDa extracellular matrix protein that is a member of the CCN (cellular communication network factor) protein family.

CTGF was identified as an endochondral ossification genetic factor that plays a crucial role in regulating various cellular functions, including proliferation, migration, adhesion, survival, differentiation and synthesis of ECM proteins during skeletal development [15]. CTGF is also involved in complex biological processes such as cell proliferation, angiogenesis, chondrogenesis, osteogenesis, wound healing, and various pathologies such as tumor development and tissue fibrosis [16, 17]. It was found that CTGF has the potential to promote tendon regeneration [18].

CTGF is a good cytokine that promotes the secretion of collagen in tenocytes, with obvious effect, and can be used as an inducer to promote tendon injury repair after tendon rupture [19, 20]. In this study, we used CTGF to induce tendon differentiation of TDSCs, and explored the signaling pathways in this process and the effect of modified human tendon stem cells (hTDSC) in promoting graft ligamentation and tendon-bone healing after ACL reconstruction.

## Materials and Methods

### Extraction of hTDSC

HTDSC was isolated and cultured. The tendon tissue of the clinical tissue sample was rinsed three times on ice with PBS containing 4% double-antibody, and the surface part of the connective tissue was cut off to separate the tendon tissue. The tissue was cut into 1 mm × 1 mm × 1 mm pieces, digested with 2 mg/mL type I collagenase at 37 °C for 2 h, and the cell suspension was collected by filtration through a 300-mesh cell sieve. The cell suspension was centrifuged at 300 × g for 10 min, the supernatant was discarded, the cell pellet was resuspended in stem cell complete medium, and seeded into a 10 cm diameter petri dish at a density of 500 cells/cm2, and culture it in a 37 °C and 5% CO2 incubator for 7 ∼ 10 days. The formation of cell colonies was observed under a microscope. The medium was changed once at 24, 48, and 72 h after primary cell inoculation, and every 3 days thereafter. The cells were subcultured after reaching 90% of the bottom of the bottle. The culture medium was discarded, washed with sterile PBS twice, digested with 0.25% trypsin for 3 ∼ 5 minutes, and the complete medium was added to terminate the digestion; centrifuge the cell suspension at 300×g for 5 min, discard the supernatant, add 2 mL of complete medium to resuspend the cells, count the density of cells at 5×10^3^ cells/cm2 seeded into cell culture dishes. The 1st ∼ 3rd generation cells were taken for subsequent experiments. The morphological changes of cells were observed by an inverted phase-contrast microscope.

### Flow characterization

Cell surface markers were detected by flow cytometry. The cells in good growth condition of P2 generation were harvested, digest with 0.25% trypsin, centrifuged at 4°C and 1000r / min for 5min. The cells were washed with PBS for 3 times, and the cells were counted. Monoclonal antibodies CD44-FITC, CD90-APC, and CD105-PE were added to each tube in turn. At the same time, negative control and two parallel tubes were set up for each tube. Incubate in dark ice for 45 min, wash the cells with PBS for 3 times to remove unbound antibody, and use 500 μL PBS resuspended cells, and flow cytometry was used for detection and analysis. Multi-lineage differentiation For differentiation studies, P2 human tendon cells were seeded in 6 wells (50,000 cells/well).

Osteogenic, chondrogenic and adipogenic differentiation were induced by applying a differentiation medium.Osteogenic Differentiation Medium Kit (Saiye Bio, RASTA-90021, CBPI0513A0); Chondrogenic Differentiation Medium Kit (Saiye Bio, MUXTA) −9004, CBPI0518A0); Adipogenic induction differentiation medium kit (Saiye Bio, RASMD-90031, CBPI0228A6). After 4 weeks, Alizarin Red, Alicia Blue and Oil Red O staining were performed to assess osteogenic, chondrogenic and adipogenic differentiation, respectively.

### Immunofluorescence staining

The induced hTDSC were fixed with 4% paraformaldehyde and washed three times with PBS for 5 min each time. Permeabilized with permeabilization solution for 10 min, washed 3 times with PBS; blocked with goat serum for 60 min at room temperature. Remove the drying blocking solution, primary antibodies COL1A1 Rabbit pAb (abclonal, A16891), COL2A1 Rabbit pAb (abclonal, A19308), COL3A1 Rabbit pAb (abclonal, A3795), HES1 (abclonal, A0925), GSK3β (abclonal, A6164), β-catenin (abclonal, A19657), TCF4 (abclonal, A1141), β-Actin (abclonal, AC026), KIT (abclonal, A0357), CAL101 (abclonal, A16891), Dcn (abclonal, A1669), TNC (abclonal, A1927)), SCX (abcam, ab58655), Myc (abclonal, AE070) were incubated at 4 °C overnight. The specimens were washed three times with PBS for 5 min each. Secondary antibody (abclonal, AS014) working solution was incubated at room temperature for 1.5 h. The specimens were washed three times with PBS for 5 min each. Nuclei staining: DAPI was added dropwise and incubated in the dark for 5 min, and the excess DAPI was washed off with PBS for 5 min×4 times. Cover slides with an anti-fluorescence quencher and store in a humidified box. The Guangzhou Mingmei upright fluorescence microscope imaging system was used to observe the conditions of each group at low magnification, and then collect the experimental results at 400X.

### Tendon induction

The experiment was divided into four groups: no induction group; induction group for 3 days; induction group for 5 days; induction group for 7 days. The isolated human tendon stem cell P2 cells were used for the experiment, and the cells in the logarithmic growth phase were selected and induced by adding a final concentration of 25 ng/mL CTGF recombinant protein. The cells were collected on 3 days, 5 days and 7 days respectively, and samples were sent for subsequent experiments.

### Lentiviral infection

After TDSC cells were cultured, HA-CTGF and Flag-KIT lentivirus were simultaneously infected with TDSC cells, and then resistance screening was performed to construct a stable overexpressing TDSC cell line. Human Flag-KIT-XbaI-F: 5’-cgTCTAGA ATGAGAGGCGCTCGCGGCGCCTGGG-3’, Human Flag-KIT-BamHI-R: 5’-cgGGATCC TCAGACATCGTCGTGCACAAGCAGA-3’.

### Co-immunoprecipitation (Co-IP, reverse pull-down validation) assay

Co-IP kit (Boxin Bio, Bes3011). Extract protein: wash the sample twice with pre-cooled 1mL 1×PBS, blot dry 1×PBS for the last time; add 1mL Cell lysis buffer and 10μL Protease inhibitors according to the amount of cells or samples; fully lyse on ice for 30min, invert and mix during 3 times. Centrifuge, 4°C, 14000g, 15min, and collect the supernatant. Incubation of protein and antibody: Take 1 / 5 volume of the above samples as input samples; Take 4 / 5 volume as IP group; store the Input sample at −20°C for later use; add 2 μg of antibody to the IP group tube; Shake and incubate in a vertical mixer at 4 °C overnight. Preparation of protein-A/G magnetic bead suspension: take 40 μL of ProteinA/G-MagBeads, stand on a magnetic stand for adsorption for 30 s, and remove the supernatant; add 500 μL of pre-cooled Cell lysis buffer, mix well, 4°C, stand on a magnetic stand for adsorption 30s, remove the supernatant; repeat the previous step twice; use 400μL of pre-cooled Cell lysis buffer to make a suspension of ProteinA/G-MagBeads. Binding of Protein-A/G magnetic beads to antibodies: Add 200 μL of the obtained ProteinA/G-MagBeads to a 1.5 mL EP tube (IP group), at 4°C, shake with a vertical mixer for 2 h; let the magnetic stand for adsorption for 30 s, go to the supernatant. Washing: let the obtained IP sample stand on a magnetic stand for adsorption for 30 s, remove the supernatant, wash the magnetic beads with 1 mL of Wash Buffer I, place it in a vertical mixer at 4°C and shake for 5 min, repeat at least 4 times; stand on the magnetic stand for adsorption For 30 s, remove the supernatant; wash once with 1 mL of Wash

Buffer II, shake with a vertical mixer at 4°C for 5 min; let it stand for adsorption on a magnetic stand for 30 s, and remove the supernatant. Elution: Add 100 μL of Elution buffer to the obtained magnetic beads of the IP group, denature and elute in a boiling water bath for 10 min, centrifuge at 4000g for 2 min at 4°C, collect the supernatant, and store at −20°C for use. Protein detection: Take 15 μL of the IP and IgG samples obtained in step 6 and the Input sample obtained in step 3, add 1/4 volume of each 5×SDS loading buffer, and bath in boiling water for 5 min. The denatured samples were stored at −80°C until use.

### Real-time PCR

Total RNA was extracted using RNAiso Plus Lysis Buffer. Reverse transcription (RT) reactions were performed using the Goldenstar™ RT6 cDNA Synthesis Kit Ver.2. Real-time PCR reactions were performed using 2 × T5 Fast qPCR Mix (SYBR Green I) kit. The reaction conditions were pre-denaturation at 95°C for 30 s, followed by 40 cycles of 95°C for 5 s, 55°C for 30 s, and 72°C for 30 s. Primer name and sequence HES1-F: GCTGGAGAAGGCGGACATTC, HES1-R: GGTCATGGCATTGATCTGGGTC; GSK3β-F: GAGAATCACTTGAATCGGGGAGGC, GSK3β-R: GATACAGAGGCTTGCTGCGC; TCF4-R: GGCCGGTTCCATACCCTGAG; Col1a1-F: GGAAGCTGGAAAACCTGGTCG, Col1a1-R GTTCGCCTTTAGCACCAGGTTG; Dcn-F: TCCTGATGACCGCGACTTCGA, Dcn-R: TGGATGTACTTATGCTCTGCCAGC; : CAGGAGAGACACTGAGGCACC; KIT-F: TGCTCCTACTGCTTCGCGTC, KIT-R: AACAATGCAGACAGAGCCGAT.

### Western Blot

Total protein was extracted using RIPA lysis buffer, and the supernatant was collected. Total protein was mixed with SDS sample buffer, denatured by boiling, separated by SDS-PAGE, transferred to PVDF membrane, and blocked with 5% skim milk for 1 hour. Incubate with the appropriate concentration of anti-HA tag antibody or primary antibody to β-Actin overnight at 4°C, followed by incubation with secondary antibody for 1 hour. Finally, the enhanced chemiluminescence (ECL) detection reagent was used to mix and evenly cover the whole film, and after 1 minute of reaction, it was placed in an exposure apparatus for exposure detection.

### Immunohistochemistry

After fixation with 4% paraformaldehyde, PBS was washed three times for 5 min each. Treated with 3% hydrogen peroxide for 15 min at room temperature; washed three times with PBS for 5 min each. The permeabilization solution was permeabilized for 10 min and washed 3 times with PBS. Goat serum was blocked at room temperature for 60 min. Remove the drying blocking solution, and incubate with primary antibodies Col1a1 (abclonal, A16891), Col2a1 (abclonal, A1560) and Col3a1 (abclonal, A3795) at 4°C overnight. The specimens were washed three times with PBS for 5 min each. Incubate with secondary antibody working solution for 1.5 h at room temperature. Washed 3 times with PBS for 5 min each. DAB color development for 10 min (protected from light). Washed twice with distilled water for 5 min. Hematoxylin counterstaining for 5 min. Washed with tap water and returned to blue. Differentiate with 1% hydrochloric acid and ethanol for 5 s. Dehydrate in 95% ethanol for 2 min. Change to fresh 95% ethanol and dehydrate for 2 min. Xylene was clear for 5 min. Change to fresh xylene and clear for 5 min. Neutral resin seal. Microscopically, the nuclei were blue. Images of the slices were collected using a Mshot MF53 inverted microscope produced by Guangzhou Mingmei Optoelectronics Technology Co., Ltd.

### Dual-luciferase reporter gene assay

According to the analysis of chromatin co-immunoprecipitation detection results, a binding site of HES1 in the GSK3β promoter region was selected, and the GSK3β promoter sequence containing the binding site of 2000bp (WT) and the mutated promoter sequence of the binding site of 2000bp (mut), and construct the corresponding luciferase reporter gene vector respectively. According to the ChIP detection results, the 2000bp sequence of the GSK3β promoter region was synthesized and cloned into the dual-luciferase reporter gene vector pGL4.11-basic, which was denoted as pGL4.11-WT; the binding site C1 in the 2000bp sequence of the GSK3β promoter region mutated and cloned into the dual-luciferase reporter gene vector pGL4.11-basic, denoted as pGL4.11-mut. Plasmid co-transfection: Add 50 μL of culture medium without antibiotics and serum, add 1 μg of plasmid DNA (mix by grouping), and mix with a gun; then add 1.6 μL of Nanofusion transfection reagent, and gently pipet with a gun Mix well, let stand at room temperature for 5-20 min, and add cells to 12-well plate. Group Lvx-NC + pGL4.11-NC; Lvx-NC + pGL4.11-WT; Lvx-HES1 + pGL4.11-NC; Lvx-HES1 + pGL4.11-WT; Lvx-HES1 + pGL4.11-mut.

### Animal acquisition and experimental grouping

There were 120 SD rats, weighing 220±20g, aged 8-10 weeks, and the experiment was started after 7 days of adaptive feeding. Provided by Chongqing Enswell Biotechnology Co., Ltd. Randomly divided into 4 groups, 30 animals/group: sham operation group; model group + PBS; model group + A gene no-load TDSC; model group + A gene overexpression TDSC. The corresponding TDSCs were transplanted according to groups, and CTGF was injected in situ for treatment, respectively. 3d, 7d, 14d, 21d, 28d, and 42d were set for observation or sampling at 6 time points for follow-up detection.

### Culture of rat tendon stem cells

TSCS rat tendon stem cells, provided by Beina Bio. When the cell density reaches 85%, discard the medium, wash twice with 2 mL of PBS, add 1 mL of trypsin solution containing 0.25%, and observe the cells under an inverted microscope. When the original adherent cells tend to be round, add 5 mL of culture medium to terminate the digestion, transfer to a 15 mL centrifuge tube, centrifuge at 1000 rpm for 5 min, discard the supernatant, add an appropriate amount of complete medium, blow evenly, inoculate at a ratio of 1:2, and culture in cells containing 5% CO2 The box continues to grow.

### ACL injury reconstruction model

Use 5% chloral hydrate (1mL/100g) for intraperitoneal injection anesthesia, make a 1cm small incision on the medial and posterior side of the ankle on the ipsilateral limb to separate the proximal end of the flexor digitorum longus tendon, and use ophthalmic forceps to pick up the proximal end of the flexor digitorum longus tendon. When the ligament is stretched, the toe of the rat will flex, which proves that the flexor digitorum longus tendon is separated. Another 0.5cm small incision is made on the back and medial side of the big toe of the foot to separate the distal end of the flexor digitorum longus tendon. When the distal end of the flexor digitorum longus tendon is raised, the toe of the rat will be flexed, the tendon will be cut off with ophthalmic scissors, the tendon will be pulled out, and then the muscle on the tendon will be separated with ophthalmic forceps to obtain a ligament of about 3 cm in length, and the prepared tendon will be soaked. Reserve in normal saline containing antibiotics.

A longitudinal incision of about 2 cm in length was made on the inside of the patella of the rat knee joint with a blade; the skin was cut layer by layer, the incision was extended, the subcutaneous tissue and joint capsule were separated, the knee joint was flexed, the ACL was fully exposed, and the ligament was cut; The bone grinding drill was marked by the normal ACL tibial and femoral attachment points, respectively, and the tibial and femoral tunnels were drilled; the flexor digitorum longus tendon was passed through the tibial and femoral bone tunnels from the outer opening of the inferior bone canal, and the graft was sutured and fixed to the proximal femur. and the periosteum or other soft tissues around the tunnel opening of the distal tibia; hemostasis with antibiotic-containing saline and adequately flushing the knee joint cavity, the patella is reset; the corresponding TDSCs are transplanted according to groups, and CTGF is injected in situ for treatment; For the first 5 days, intraperitoneal injection of penicillin (5U/kg) was given to fight infection.

### Behavioral observation score (Lequesne MG score)

Lequesne MG behavioral scores were performed on the 3rd, 7th, 14th, 21st, 28th, and 42nd days of modeling. The scoring indicators were divided into three categories: Local pain stimulation response, graded as I: no abnormal pain response; II: no abnormal pain response; Limb contraction; Grade III: the affected limb is contracted and spasm, accompanied by mild systemic reactions, such as trembling all over the body, turning back and sucking; Grade IV: the affected limb is severely contracted and spasm, and the whole body is trembling, running around, and struggling. The corresponding scores are 0, 1, 2, and 3. Changes in gait, graded as grade I: no limp, normal running, and strong kicking on the ground; grade II: mild lameness when running, but strong kicking on the ground; grade III: the affected limb participates in walking, but lameness Details; Grade IV: The affected limb cannot participate in walking, and cannot touch or push on the ground. The corresponding scores are 0, 1, 2, and 3. 3. Joint activity (0° when straightened): Grade I: more than 90°; Grade II: 45°∼90°; Grade III: 15°∼45°; Grade IV: 15°. The corresponding scores are 0, 1, 2, and 3. Joint swelling, grade I: no swelling, bony markers are clearly visible; grade II: mild swelling, bony markers become shallow; grade III: obvious swelling, bony markers disappear. The corresponding scores are 0, 1, and 2.

### Micro CT inspection

The specimens were placed in 4% paraformaldehyde to fix for 24 hours, rinsed properly with PBS, removed the excess muscle and ligament tissue around the knee joint, and stored the specimens in 70% alcohol at −20° for inspection. Each sample was scanned with the Scanner software of Skyscan1276 Micro CT. The tibia and femur were aligned based on the punched area, and the 0.2 mm diameter annular area around the hole was set as a three-dimensional reconstruction region of interest (ROI). CT-AN software was used to detect bone mineral density and measure the width of the bone tunnel.

### HE staining

After getting the samples, they were washed with PBS and fixed with 4% paraformaldehyde. Dehydration: soak the tissue in 75% ethanol, 85% ethanol, 95% ethanol I, 95% ethanol II, 100% ethanol I, and 100% ethanol II for 2 h each. Transparency: soak the dehydrated tissue in ½ xylene, xylene I, and xylene II for 10 min, respectively. Permeabilization: Immerse the transparent treated tissue in melted paraffin for 3h. embedded. Sectioning: Use a microtome to cut the tissue in the paraffin block into 2.5um thick slices and lay them flat on a detachment-proof glass slide. Baked slices: Place the slices on a 55°C slicer, so that the tissue slices are tightly attached to the anti-detachment glass slides. Dewaxing: soak the paraffin sections in xylene I, xylene II and 1/2 xylene, respectively, for 10 min each. Rehydration: soak the deparaffinized paraffin sections in 100% ethanol I, 100% ethanol II, 95% ethanol I, 95% ethanol II, 85% ethanol, and 75% ethanol for 5 min each. Wash with double distilled water 3 times, 2 min each time. Stain with hematoxylin for 5 min. Rinse excess staining solution with tap water for about 5 min. Wash again with distilled water (a few seconds is enough). Destain with 1% hydrochloric acid alcohol to remove excess hematoxylin staining solution in the cytoplasm. Eosin staining solution for 2 min. Rinse excess dye solution with tap water for about 1 min. Wash again with distilled water. Dehydrate in 95% ethanol for 2 min. Change to fresh 95% ethanol and dehydrate for 2 min. Xylene was clear for 5 min. Change to fresh xylene and clear for 5 min. Neutral resin seal. Microscopically, the nucleus is blue and the cytoplasm is red or pink. The images of the slices were collected using a Mshot MF53 microscope produced by Guangzhou Mingmei Optoelectronics Technology Co., Ltd.

### Transmission electron microscope observation

Prefixed with a 3% glutaraldehyde, then the tissue was postfixed in 1% osmium tetroxide, dehydrated in series acetone, infiltrated in Epox 812 for a longer, and embedded. The semithin sections were stained with methylene blue and Ultrathin sections were cut with diamond knife, stained with uranyl acetate and lead citrate. Sections were examined with JEM-1400-FLASH Transmission Electron Microscope.

### Statistical Analysis

The data in this paper are presented as mean ± SD (standard deviation). Statistical analysis was performed using GraphPad Prism 8.0 software (San Diego, CA, USA) and data significance was analyzed by t-test or two-way analysis of variance (ANOVA).

## Result

### HTDSC exhibit multi-directional differentiation potential

Morphological observation showed that the nucleated cells obtained from tendon tissue digestion were seeded at a density of 500 cells/cm^2^, and the cells grew clone-like. On 7-10 days of primary culture, many scattered clustered cell colonies were seen, which showed adherent growth from the center to the outside, and the cells were irregular in shape. The nucleus gathers in the center, and the protrusions formed by the cytoplasm are radial outwards. After the second passage cells grew to 90% confluence, the cells were uniform in shape, showing elongated spindle shape and fibroblast-like growth. (Fig. 1A)

**Fig 1.**
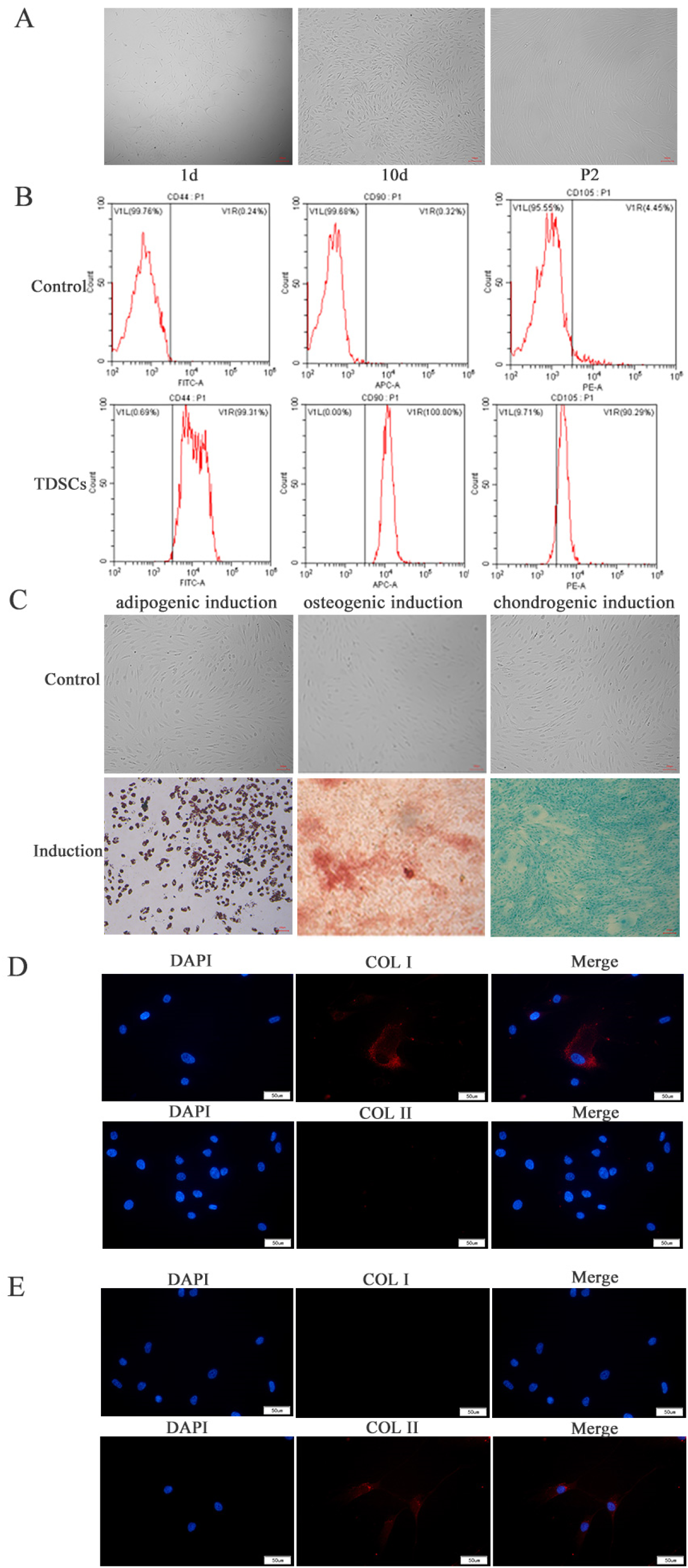
The extraction of hTDSC and the verification of their multi-directional differentiation potential. (A) Culture and passage of hTDSC; scale: 100 μm. (B) Identification of surface markers; V1L: the proportion of negative cells refers to the proportion of cells with no specific antigen markers detected; V1R: the proportion of positive cells, the proportion of cells with specific antigen markers detected by flow cytometry. (C) Oil red O staining to verify the adipogenic induction of hTDSC, Alizarin red staining to verify the osteogenic induction of hTDSC, and Alicia blue staining to verify the chondrogenic induction of hTDSC; Scale: 100 μm. (D) Expression of type I and type II collagen in hTDSC after osteogenic induction and (E) chondrogenic induction; Scale: 50 μm.

The specific surface markers CD44, CD90 and CD105 of hTDSCs were detected, and the positive rates were 99.24% ± 0.15%, 99.52% ± 1.52% and 89.81% ± 2.7%, respectively (Figure 1B). After adipogenic induction of P2 generation TDSCs, red lipid droplets appeared in cells stained with Oil Red O (Fig. 1C). After osteogenic induction, red deposits (i.e., calcium nodules) appeared upon alizarin red staining (Fig. 1C), and the important marker type I collagen was significantly expressed (Fig. 1D). The induced chondrocytes appeared dark blue after staining with alician blue (Fig. 1C), and the chondrogenesis marker type II collagen was significantly expressed (Fig. 1E).

The primary isolated hTDSC have uniform morphology, stable growth, positive cell surface markers CD44, CD90 and CD105, and can undergo tri-lineage differentiation under corresponding induction conditions, with typical stem cell characteristics.

### Expression of DEGs during the differentiation of tendon stem cells into tendon

The cells in the logarithmic growth phase of the P2 generation were selected, and the final concentration of 25 ng/mL CTGF recombinant protein was added to induce tendon formation of TDSCs. The cells were collected on days 0, 3, 5, and 7 and sent for single-cell sequencing. A total of 14 cell types were identified in the 4 groups of samples sequenced (Fig. 2A), and the main cell groups were annotated, namely Tenocytes (tenocytes), Tendon derived stem cells (tendon derived stem cells), and Smooth muscle cells (smooth muscle cells). cells), mesenchymal stem cells (mesenchymal stem cells) (Fig. 2B). Compared with the non-induction group, the number of tenocytes in the 3-day induction group showed an increasing trend, but the overall cell population did not change significantly; The number of stem cells decreased significantly, and the distribution of cell populations was significantly changed; there was no significant change between the 7-day induction group and the 5-day induction group (Fig. 2C, D). The above results indicated that the expression of proteins in the cells had changed significantly within 5 days after the tendon-derived stem cells were induced by CTGF. The screening of tendon differentiation-related proteins can be carried out on the differentially expressed genes between 0 days, 3 days and 5 days of induction of tendon-derived stem cells.

**Fig 2.**
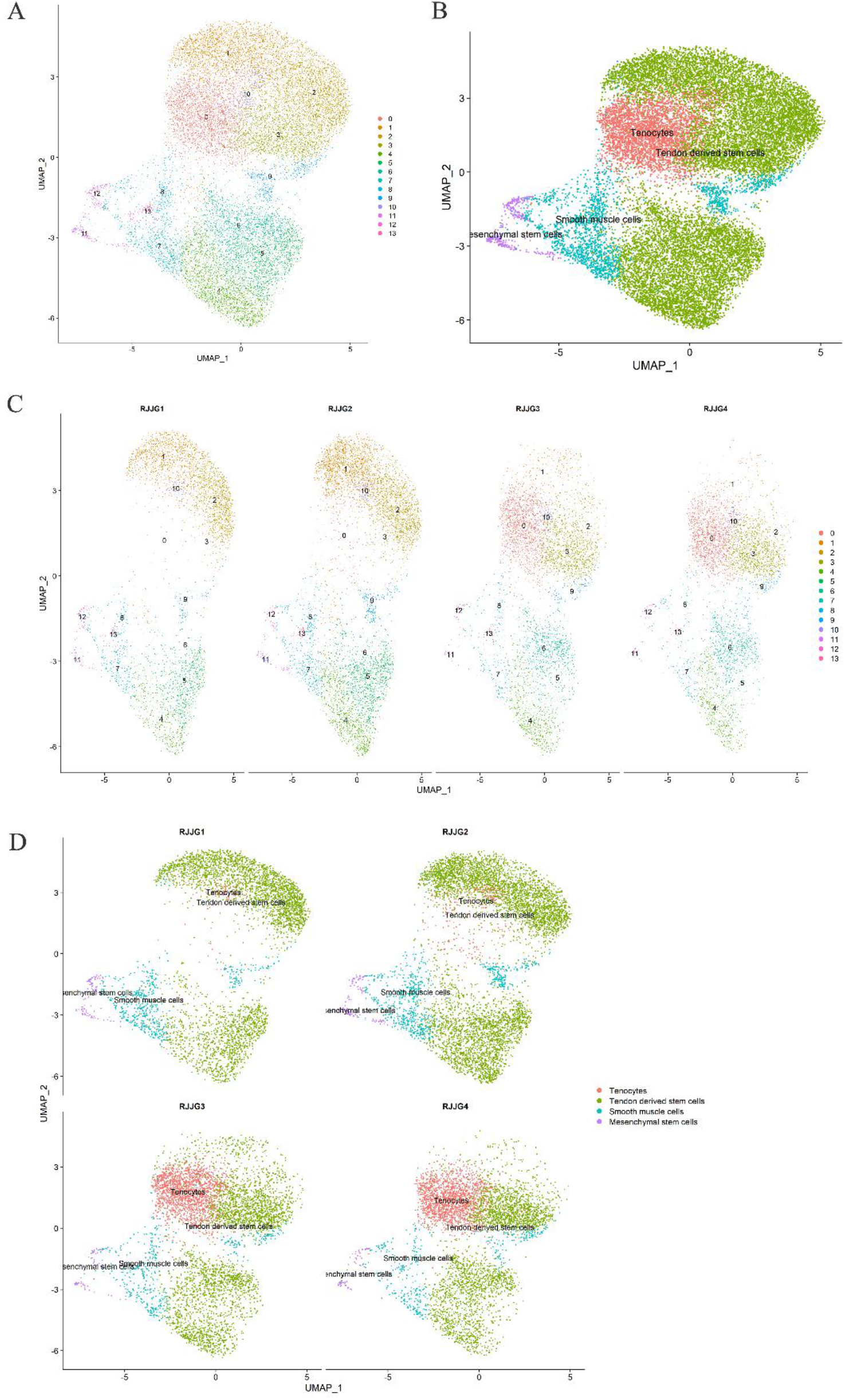
Induced cell morphology and single-cell sequencing analysis. (A)Cell clustering cluster map, (B)Cell clustering cluster annotation map, (C)Clustering diagram of each group of cells, (D)Clustering annotation map of each group of cells. RJJG1, RJJG2, RJJG3, RJJG4, represent the non-induction group, the induction group for 3 days, the induction group for 5 days and the induction group for 7 days, respectively.

According to the results of GO and KEGG analysis, most of the enriched signaling pathways of differentially expressed genes between the 0-day, 3-day, and 5-day induction groups have little correlation with the suppressed differentiation-related pathways (Fig.3 and Fig. 4). However, through GO analysis, we can see that translational initiation, ATP metabolic process, cadherin binding, etc are highly enriched in terms of function, so there is an undetected or yet unexplained signaling pathway in the cells associated with tendon differentiation (Fig.3).

**Fig 3.**
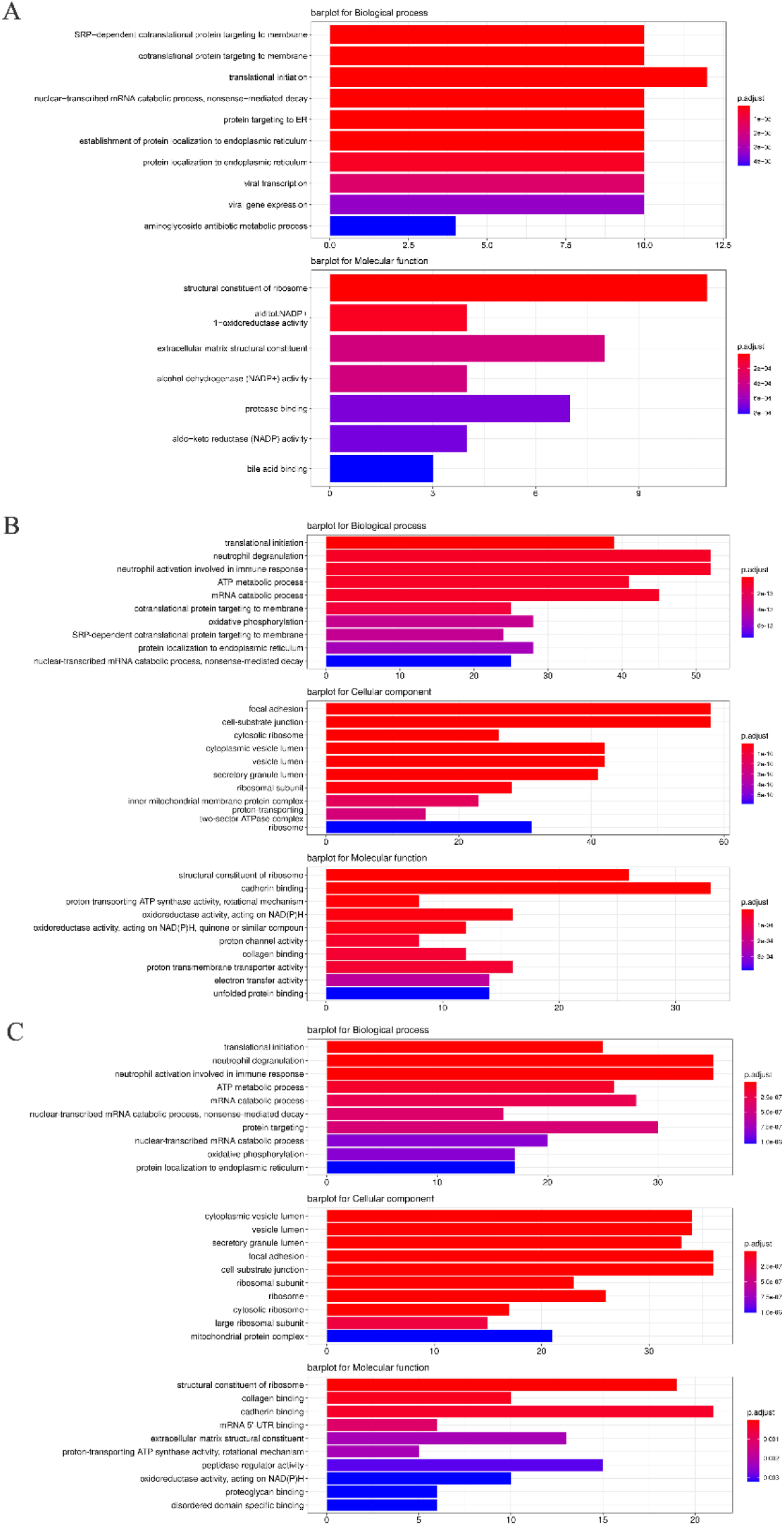
GO-TERM analysis, (A) The GO analysis of the DEGs of the 0-day group and the 3-day group; (B) The GO analysis of the DEGs of the 0-day group and the 5-day group; (C) The GO analysis of the DEGs of the 3-day group and the 5-day group.

**Fig 4.**
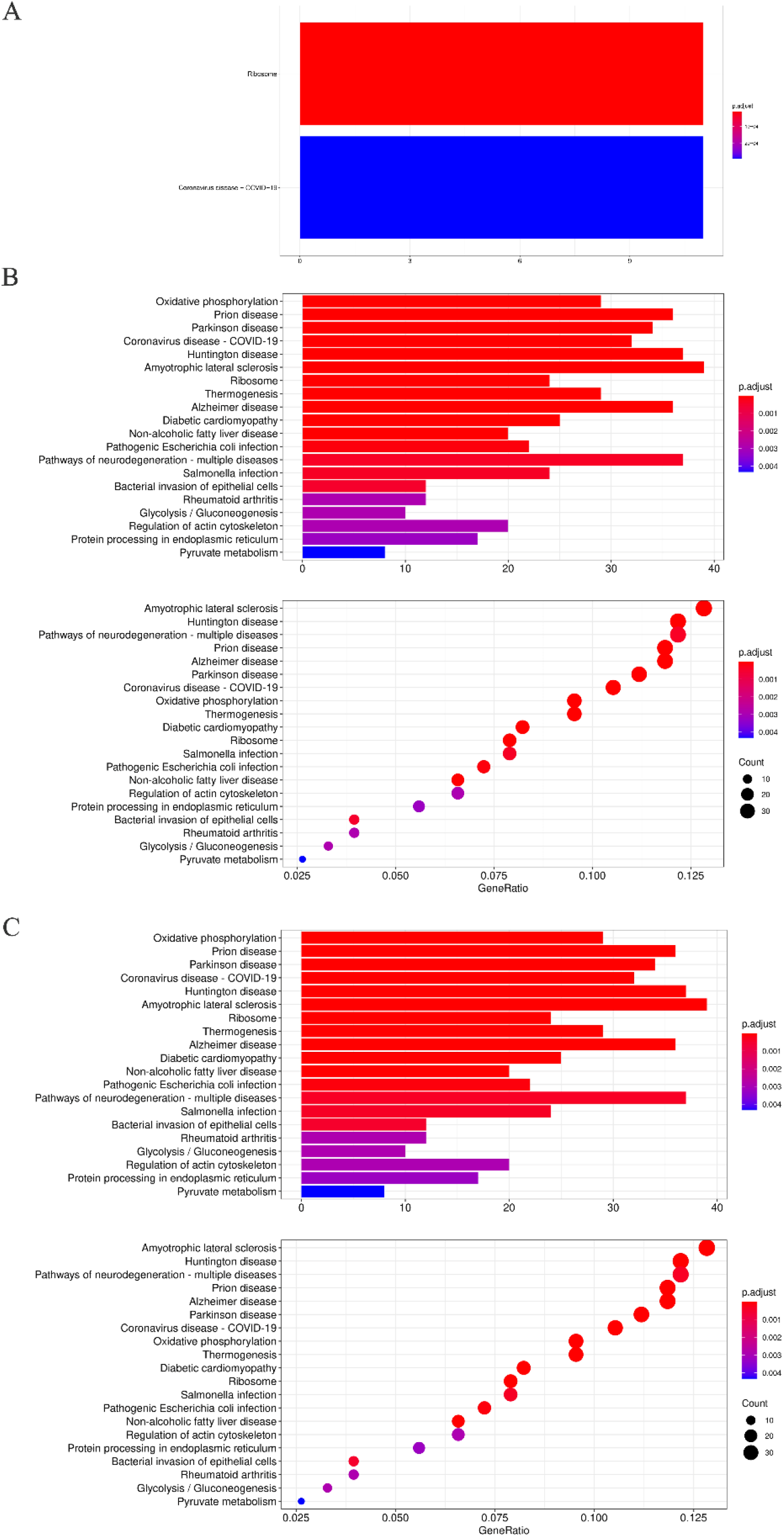
KEGG analysis. (A) KEGG analysis of DEGs of the 0-day group and 3-day group; (B) KEGG analysis of DEGs of the 0-day group and 5-day group; (C) KEGG analysis of DEGs of the 3-day group and 5-day group.

### CTGF and Kit have direct interactions

After human HA-CTGF overexpressing lentivirus infected TDSCs, co-immunoprecipitation (Co-IP) was performed. The results showed that HA-CTGF protein was detected in the Input group and IP group, but not in the IgG group, indicating that the pull-down product contained HA-CTGF protein, and the IP-grade antibody was specifically bound to HA-CTGF (Fig. 5B). After the mass spectrometry data retrieval, PSM FDR ≤ 0.01 and Protein FDR ≤ 0.01 were screening criteria (Fig. 5A). The statistical results of the number of proteins and peptides identified by the final sample are shown in Table 1. According to the results of single-cell sequencing results and mass spectrometry, it is estimated that the expected pathway: CTGF interacts with membrane receptor protein KIT (mutual protein KIT) to regulate candidate gene HES1 (transcription factor HES1) to regulate Wnt / β-catenin signaling pathway (GSK3β, β-catenin, TCF4).

**Fig 5.**
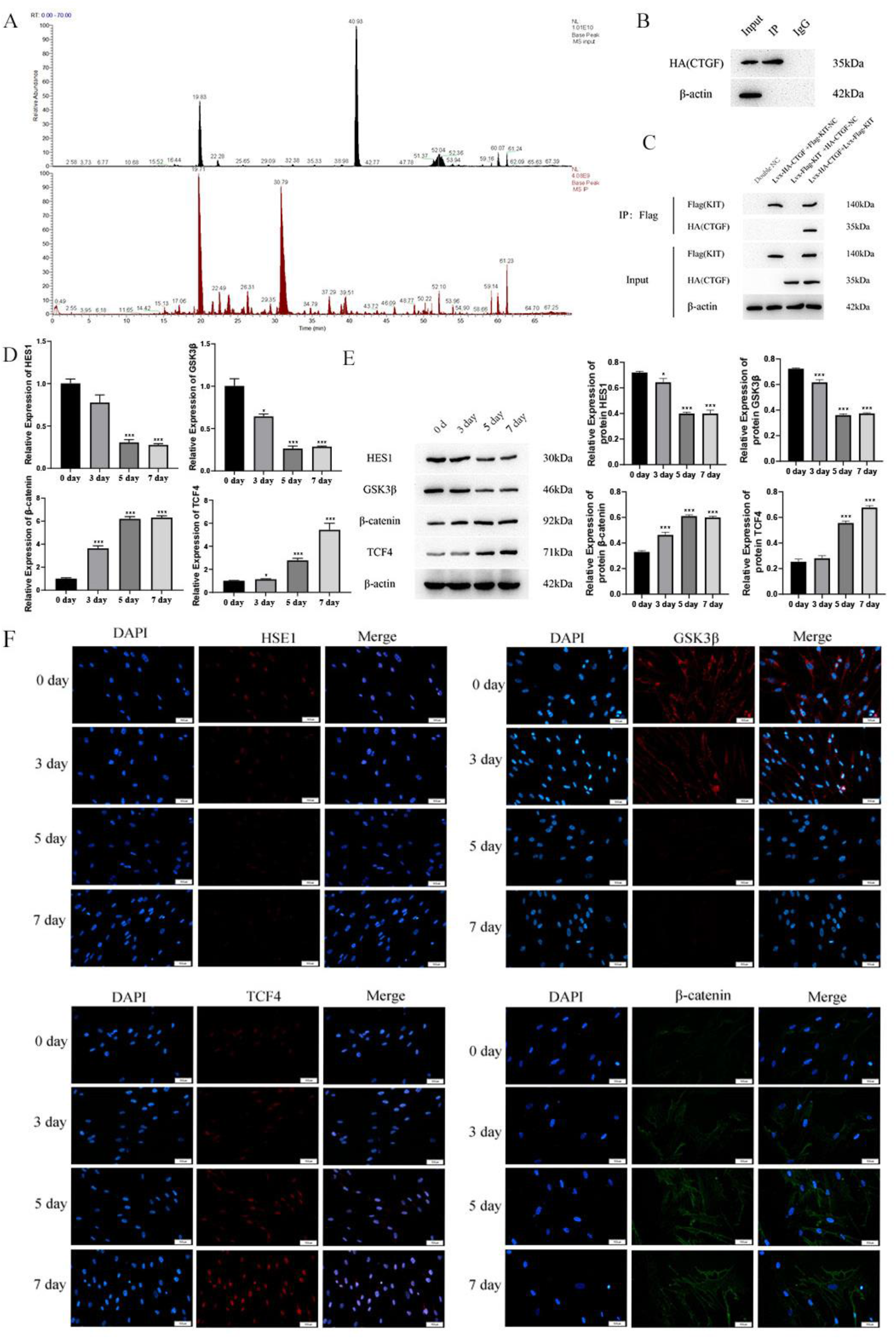
(A) Mass spectrum Basepeak of Input and IP mass spectrometry samples; (B) Expression of CTGF protein in Input group, IP group and IgG group; (C) The expression of KIT and CTGF proteins; The mRNA and protein expressions of HES1, GSK3β, β-catenin and TCF4 were detected by qPCR(D), WB (E) and immunofluorescence (F).Note: Double NC: double no-load group; Lvx-Flag-KIT+ HA-CTGF-NC: Flag-KIT overexpression group+HA-CTGF no-load group; Flag-KIT-NC+ Lvx-HA-CTGF: HA-CTGF overexpression+Flag-KIT no-load group; Lvx-Flag-KIT+ Lvx-HA-CTGF: HA-CTGF overexpression+Flag-KIT overexpression group. “*” means p<0.05 compared with the non-induction group, “**” means p<0.01 compared with the non-induction group, “***” means p<0.001 compared with the non-induction group, the significant difference in the figure Sex was compared to the non-induced group.Scale: 50 μm

The preliminary verification of speculative pathways was made with reverse Co-IP, QPCR, WB, and immunofluorescence. The results show that only when both Flag-KIT and HA-CTGF are expressed, the expression of Flag-Kit and HA-CTGF can be detected in the drop-down product, which proves that there is a direct interaction between KIT and CTGF (Fig. 5C).

The mRNA and protein expression levels of tendon markers [21] HES1, GSK3β, β-catenin, and TCF4 were measured in four groups: non-induction group, induction group for 3 days, induction group for 5 days, and induction group for 7 days. Compared with the non-induction group, the expression levels of HES1 and GSK3β in the induction group gradually decreased, while the expression levels of β-catenin and TCF4 increased with the increase of induction days. The expression levels of HES1, GSK3β and β-catenin in the 5-day induction group and the 7-day induction group were similar (Fig. 5D, E). The protein expression levels of HES1, GSK3β, β-catenin and TCF4 in the four groups by immunofluorescence were consistent with the results of qPCR (Fig.5F). In conclusion, there is a direct interaction between CTGF and KIT, and the expression levels of the screened indicators are consistent with the single-cell sequencing results, or the expression changes are in line with expectations.

### KIT correlates with predicted signaling pathways and stem cell differentiation

It was proved by gene regulation that the expression change of the KIT gene was positively correlated with the expression of tendon markers Col1a1, Dcn, TNC, Scx and the tendon differentiation of stem cells.

The changes of HES1, GSK3β, β-catenin, and TCF4 with KIT gene expression are consistent with the expected changes, and from the changing trend, GSK3β is likely to be a downstream target gene regulated by HES1 transcription (Fig. 6 and Fig. 7). The expression of collagen proteins was detected by immunohistochemistry. The results showed that Col1a1 and Col3a1 were highly expressed in the KIT overexpression group but decreased in the KIT interference group. There was no significant change in the expression of Col2a1 (Fig. 8).

**Fig 6.**
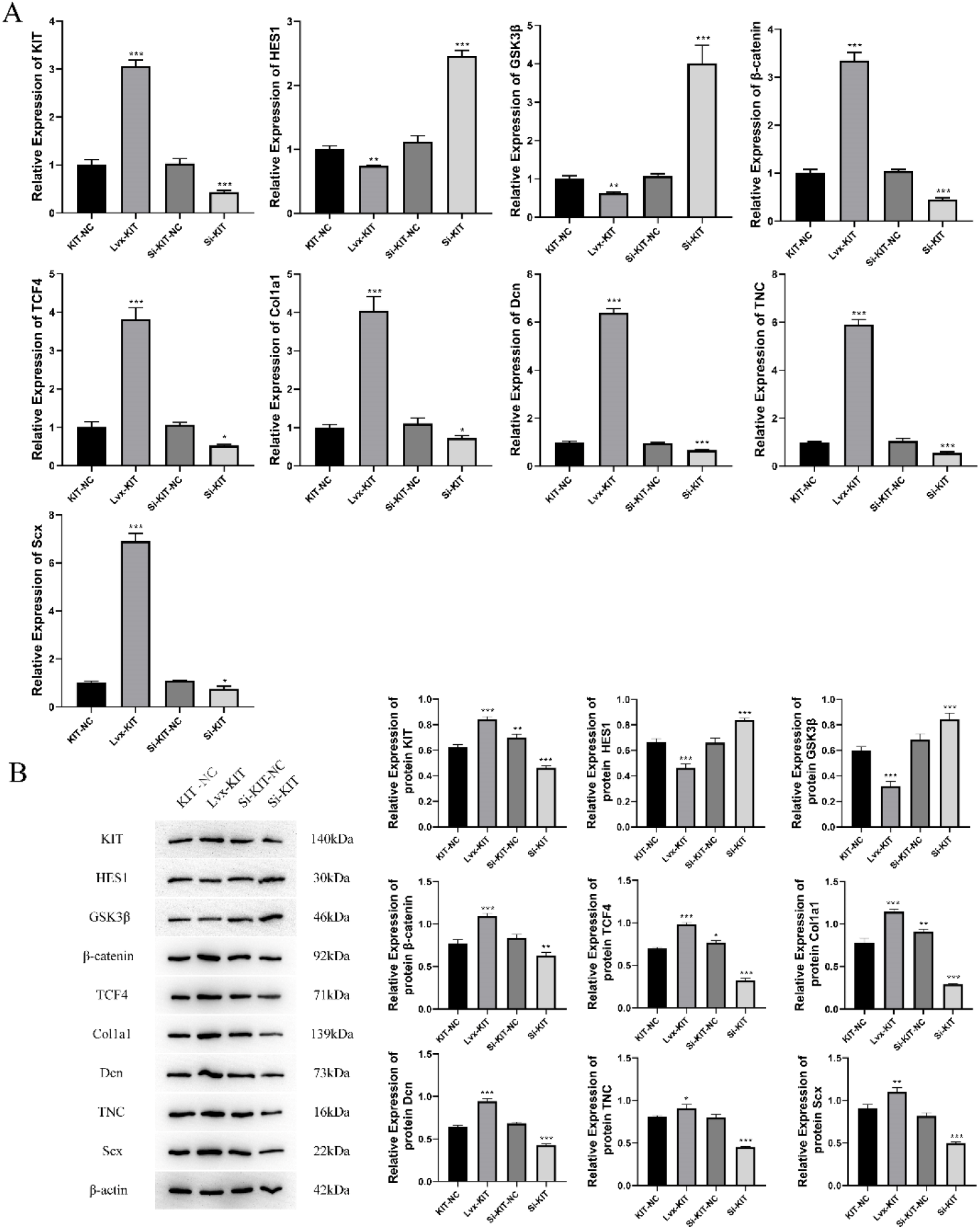
The expression levels of KIT, HES1, GSK3β, β-catenin, TCF4, Col1a1, Dcn, TNC, Scx mRNA and protein were detected by qPCR (A) and WB (B). Note: KIT-NC: KIT no-load group; Lvx-KIT: KIT overexpression group; si-KIT-NC: KIT interference no-load group; si-KIT: KIT interference group. “*” means p<0.05, “**” means p<0.01, “***” means p<0.001, the significant difference in the figure Sex was compared to the KIT-NC group.

**Fig 7.**
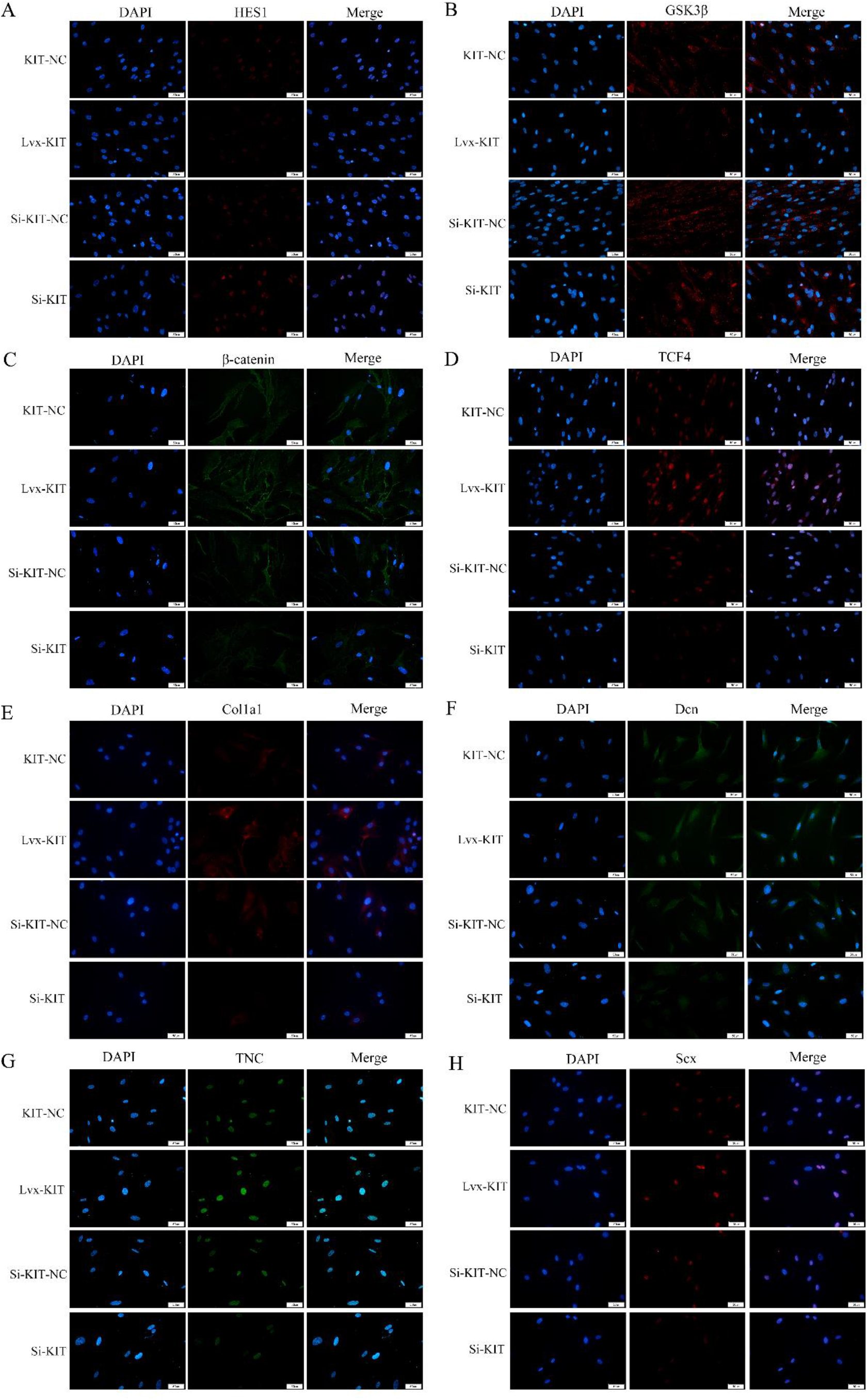
Immunofluorescence detection of protein expression levels of HES1, GSK3β, β-catenin, TCF4, Col1a1, Dcn, TNC and Scx. Scale: 50 μm.

**Fig 8.**
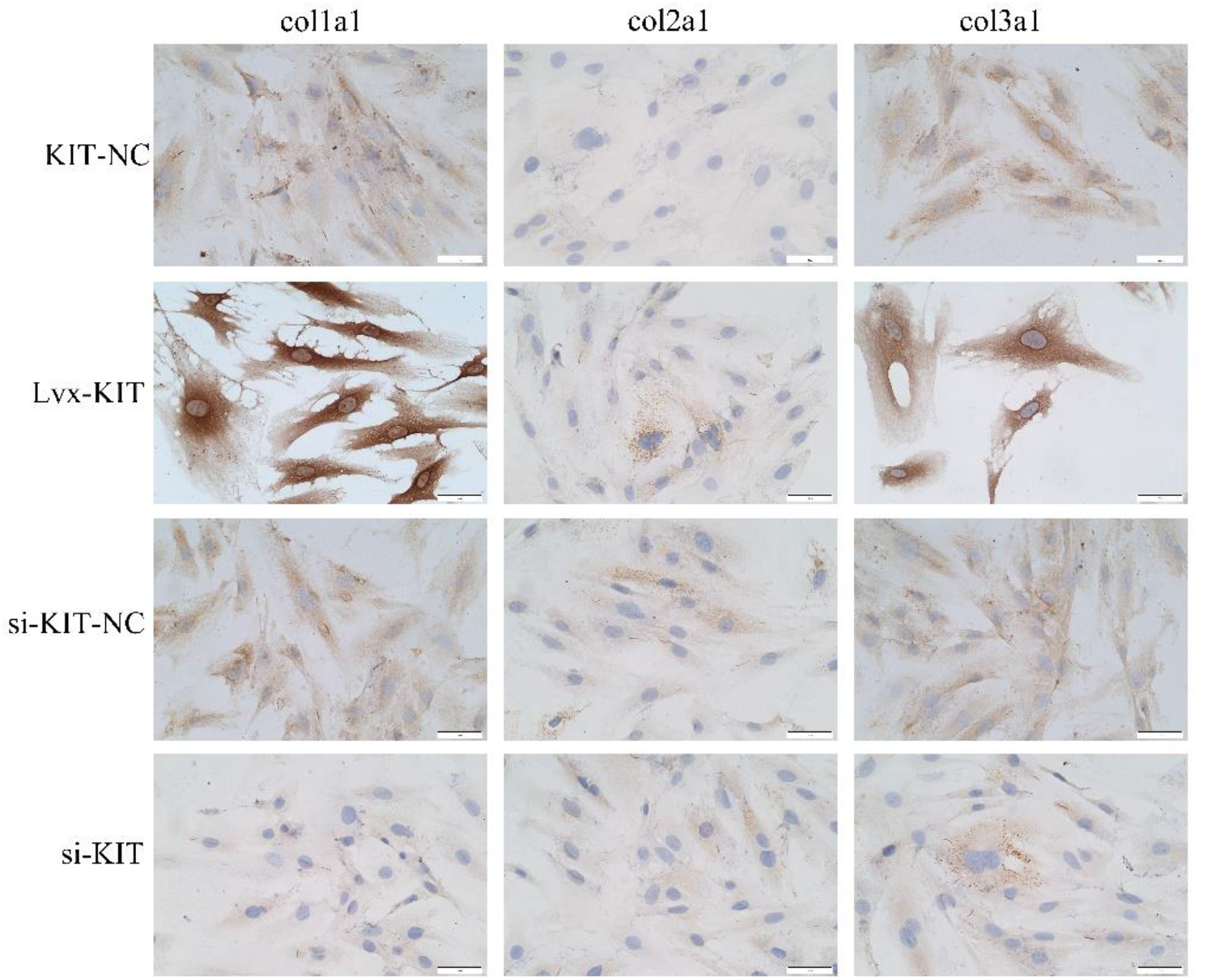
Expression levels of collagen Col1a1, Col2a1 and Col3a1 were detected by immunohistochemistry. Note: KIT-NC: KIT no-load group; Lvx-KIT: KIT overexpression group; si-KIT-NC: KIT interference no-load group; si-KIT: KIT interference group.Scale: 50 μm.

By interfering with the expression of KIT gene to detect the changes of related indicators, and then overexpressing KIT gene based on the interfering KIT gene (that is, restoring) and then detecting the changes of related indicators. It was found that the restoration of KIT gene expression corresponds to the change of predicted signaling pathway proteins and stem cell differentiation, proving the role of KIT gene and predicted signaling pathway in stem cell differentiation (Fig.9, 10, 11).

**Fig 9.**
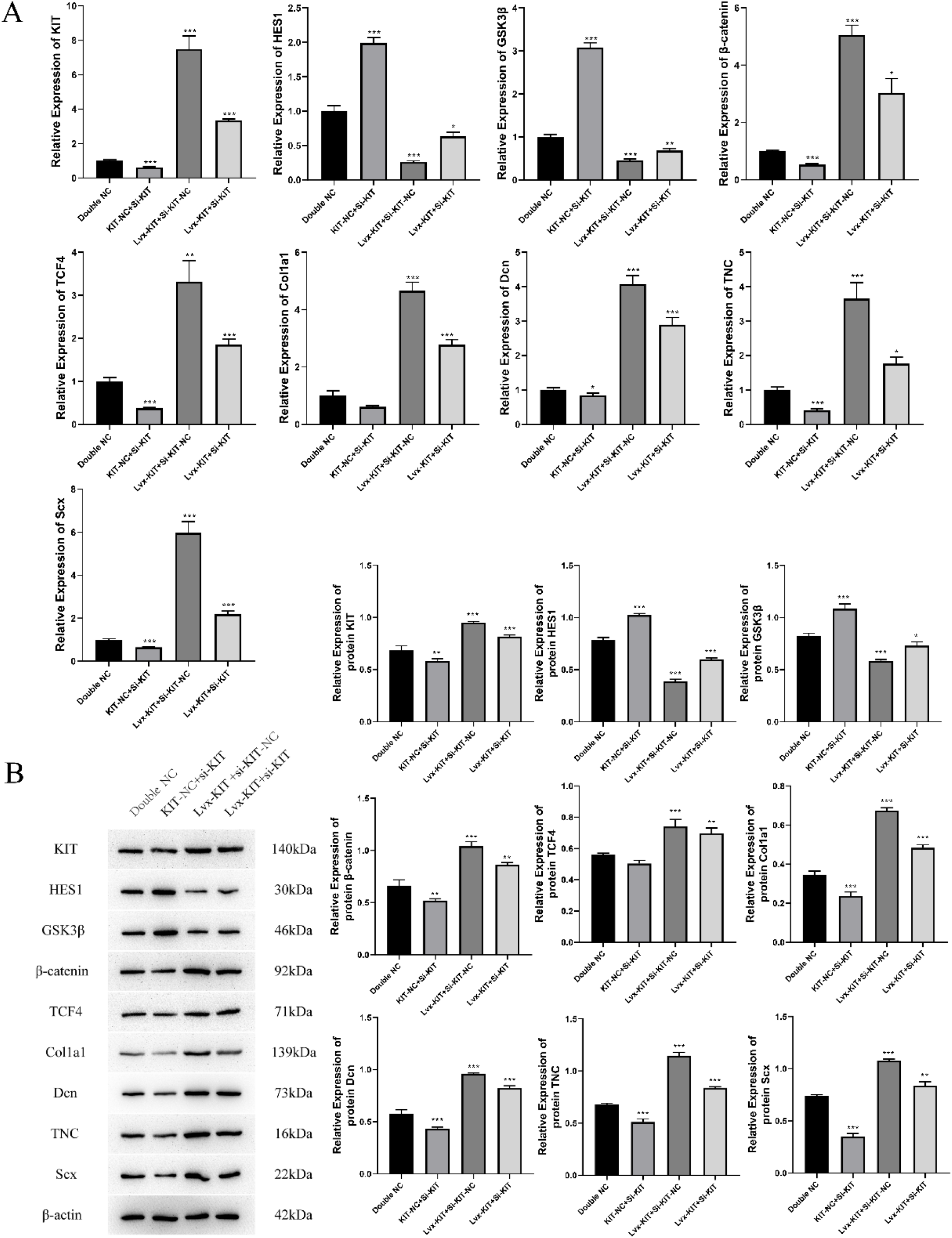
The expression levels of KIT, HES1, GSK3β, β-catenin, TCF4, Col1a1, Dcn, TNC, Scx mRNA and protein were detected by qPCR (A) and WB (B). Note: Double NC: double no-load group; KIT-NC+si-KIT:KIT no-load group+ KIT interference group; Lvx-KIT+si-KIT-NC: KIT overexpression+ KIT interference no-load group; Lvx-KIT+ si-KIT: KIT overexpression + KIT interference group. “*” means p<0.05, “**” means p<0.01, and “***” means p<0.001, the significant difference in the figure Sex was compared to the Double NC group.

**Fig 10.**
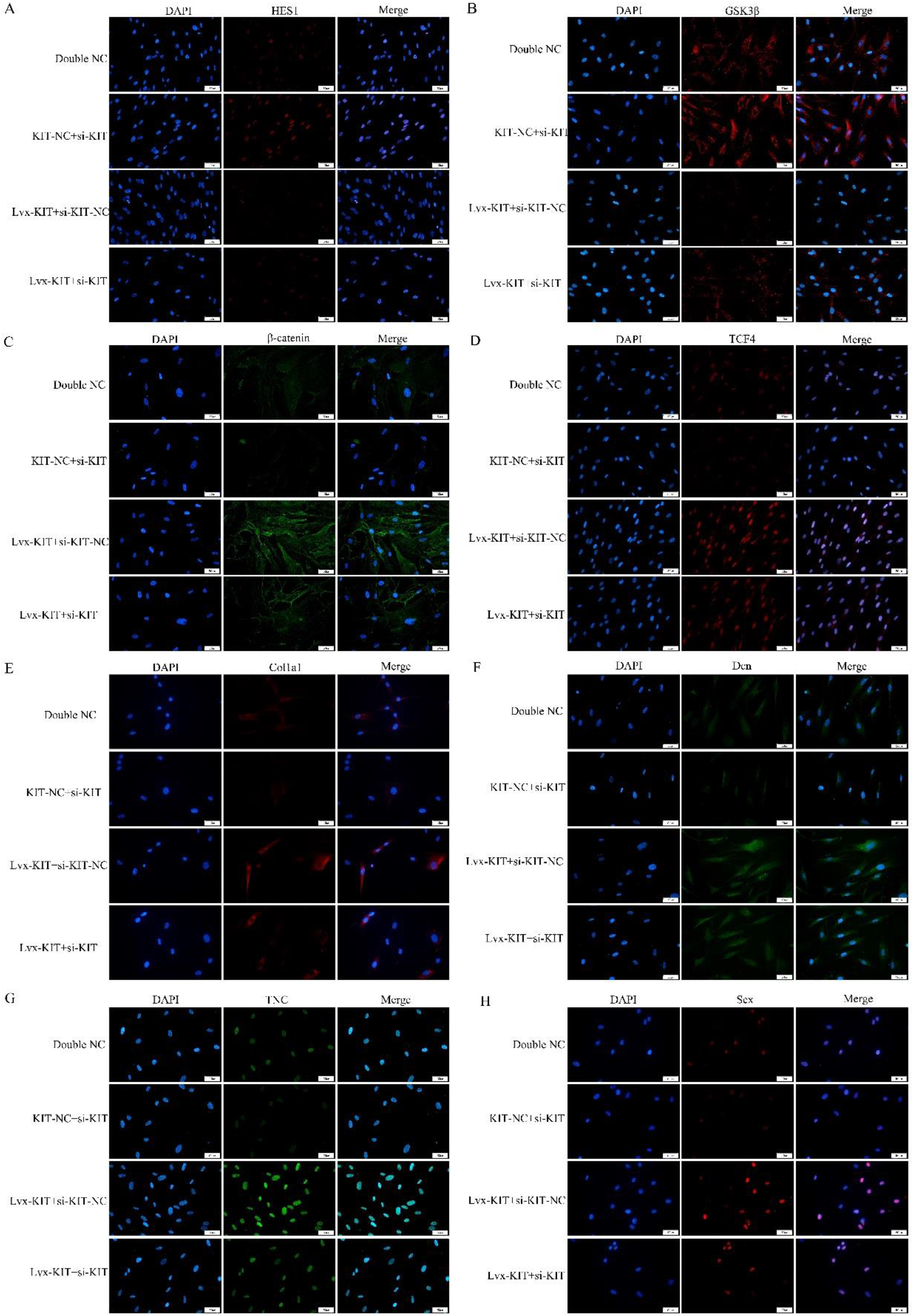
Immunofluorescence detection of protein expression levels of HES1, GSK3β, β-catenin, TCF4, Col1a1, Dcn, TNC and Scx.Note: Double NC: double no-load group; KIT-NC+si-KIT:KIT no-load group+ KIT interference group; Lvx-KIT+si-KIT-NC: KIT overexpression+ KIT interference no-load group; Lvx-KIT+ si-KIT: KIT overexpression + KIT interference group. Scale: 50 μm.

**Fig 11.**
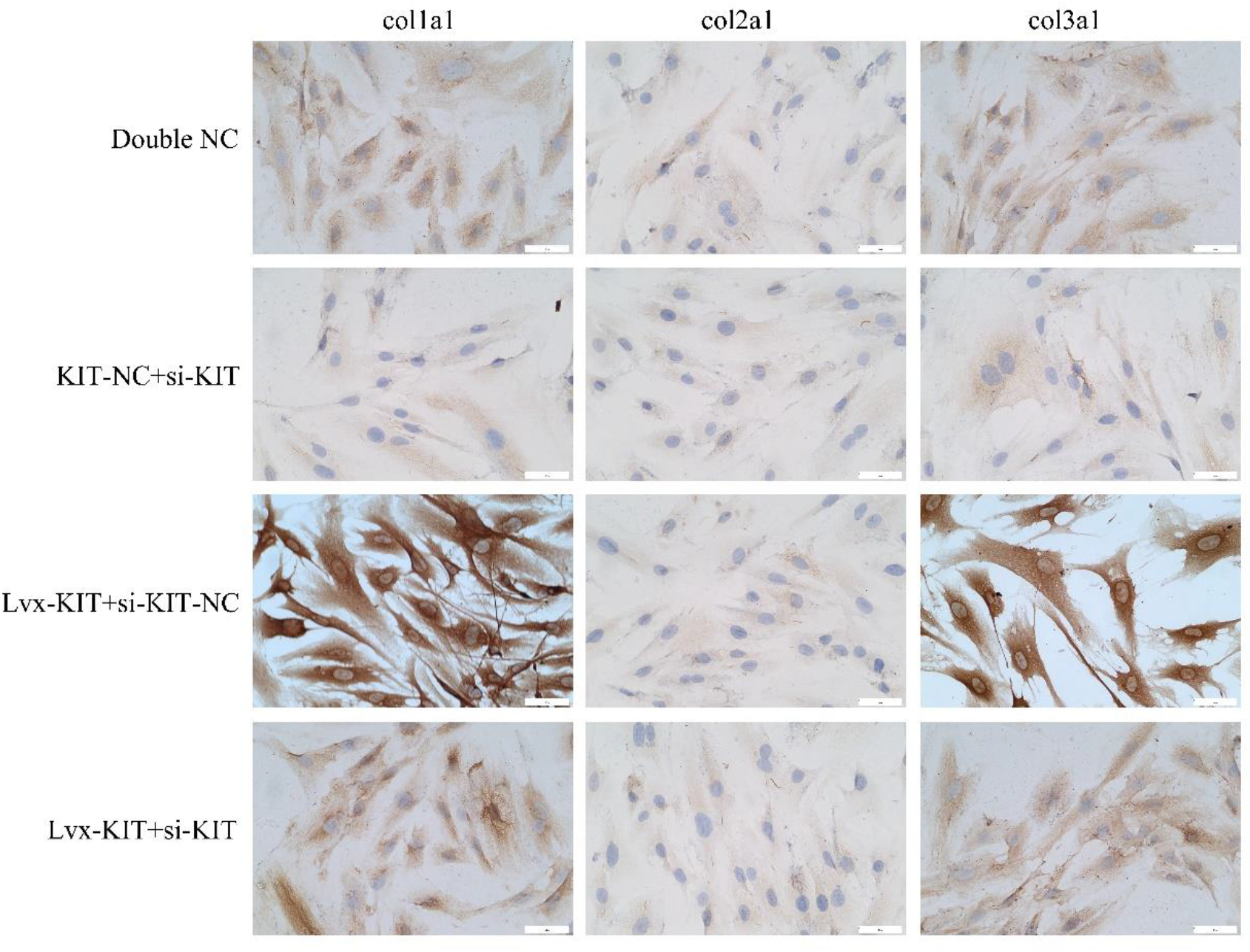
Expression levels of collagen Col1a1, Col2a1 and Col3a1 were detected by immunohistochemistry. Note: Double NC: double no-load group; KIT-NC+si-KIT:KIT no-load group+ KIT interference group; Lvx-KIT+si-KIT-NC: KIT overexpression+ KIT interference no-load group; Lvx-KIT+ si-KIT: KIT overexpression + KIT interference group. Scale: 50 μm.

### GSK3β is a target gene of the transcription factor HES1

ChIP experiments confirmed that the Myc tag was detected in both the IP group and the Input group, indicating that the pull-down product was a Myc-tag antibody-specific pull-down product (Fig. 12A). The JASPAR2020 database was used to predict the binding site of the transcription factor HES1 to GSK3β, β-catenin, and TCF4 gene promoter region 2000bp sequence. Three binding sites (GSK3β-1∼GSK3β-3) in the GSK3β promoter region were obtained, which are marked as C1∼C3; 1 binding site in the β-catenin promoter region (β-catenin-1), denoted as D1; 3 binding sites in the TCF4 promoter region (TCF4-1∼TCF4-3), denoted as E1∼E3.

**Fig 12.**
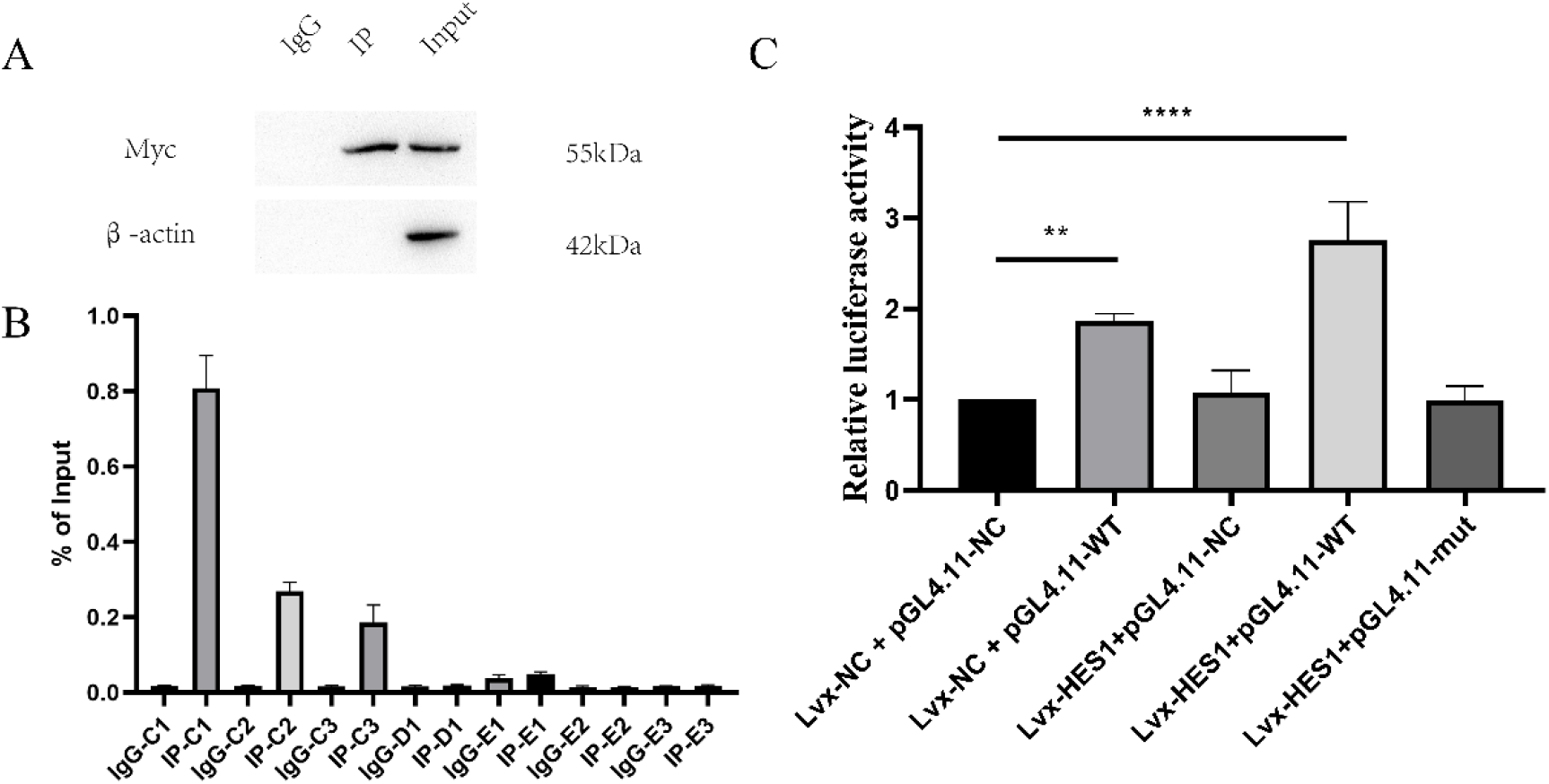
Validation of transcription factor HES1 target genes. (A) ChIP pull-down product assay; (B) ChIP-qPCR and (C) dual-luciferase reporter assay confirmed that the transcription factor HES1 target gene was GSK3β. Note: “**” means p<0.01, “***” means p<0.001.

ChIP-qPCR detection results showed that only the binding site C1 of the GSK3β promoter sequence in GSK3β, β-catenin, and TCF4 was detected in the pull-down product (Fig. 12B). Because C1 was detected in the largest amount in the pull-down product, GSK3β was most likely the downstream target gene of the transcription factor HES1, and the binding site was C1.

The results of ChIP-qPCR were reversely demonstrated by the dual-luciferase reporter assay. The dual-luciferase reporter gene assay showed that in the presence of binding site C1 (i.e., pGL4.11-WT), the fluorescence intensity was significantly enhanced, and it increased with the increase of HES1 expression. This indicates that GSK3β is the target gene of HES1, and the binding site of HES1 in the GSK3β promoter region is C1 (Fig. 12C). The results of ChIP-qPCR and dual-luciferase reporter gene assay both proved that GSK3β is a downstream target gene regulated by HES1 transcription.

### Establishment of an animal model of ACL injury and reconstruction in rats

The mRNA and protein expressions of tendon markers and transcription factors Col1a1, Dcn, TNC, and Scx detected by qPCR and WB were significantly increased in the KIT gene overexpression group, and the expression was significantly decreased in the case of KIT interference (Fig.13A, B). The mRNA and protein expressions of the expected pathway proteins HES1, GSK3β, β-catenin and TCF4 were detected as expected (Fig.13A, B). Immunofluorescence experiments confirmed the above results (Fig.14). Immunohistochemical detection showed that the collagen proteins Col1a1 and Col3a1 were significantly increased in the KIT overexpression group and decreased in the KIT interference group. The expression of Col2a1 did not change significantly (Fig.15). In conclusion, the KIT gene-overexpressing rat TDSC stable transgenic strain and the KIT gene-interfering rat TDSC stable transgenic strain was successfully constructed, which can be induced by CTGF for tendon differentiation.

**Fig 13.**
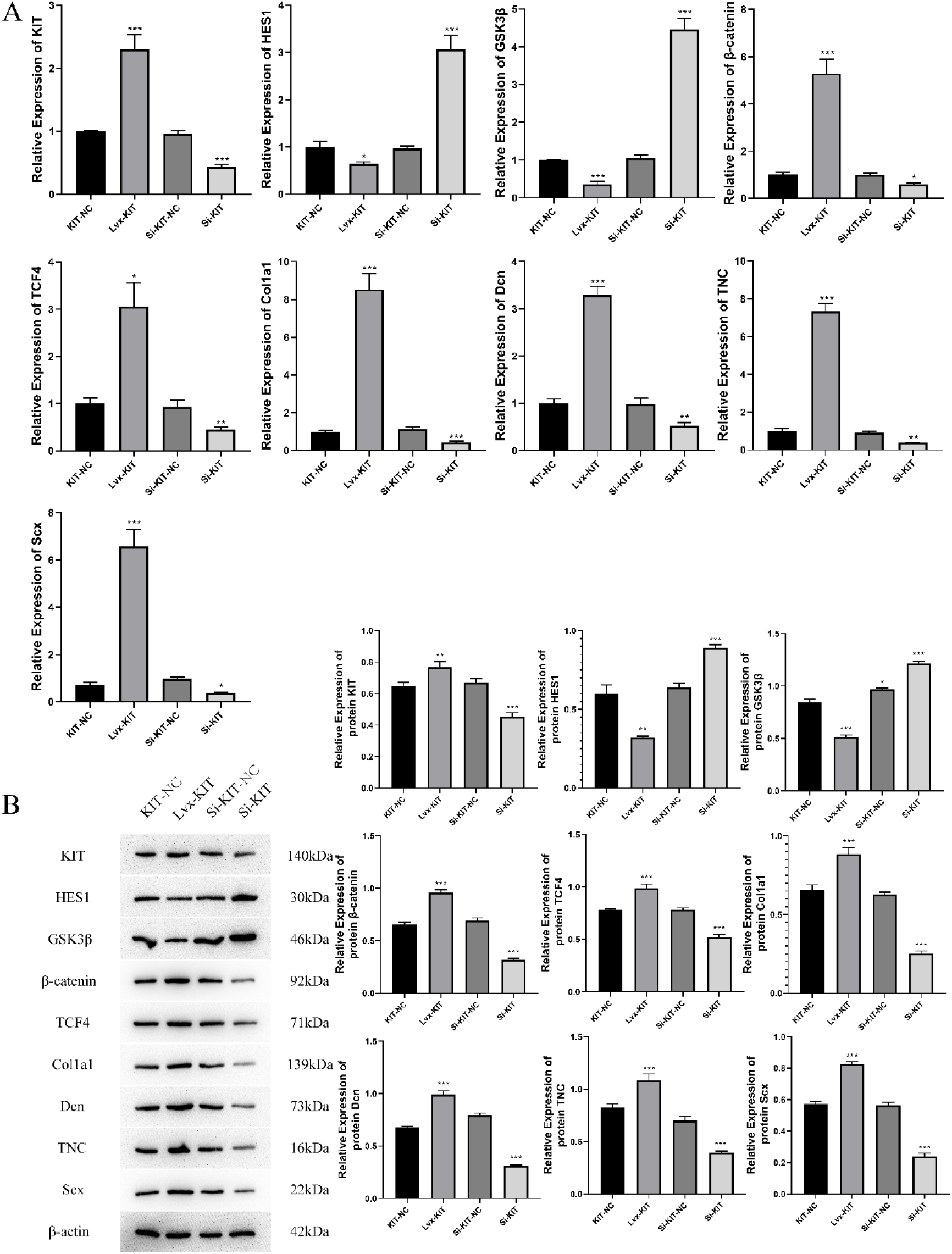
The expression levels of KIT, HES1, GSK3β, β-catenin, TCF4, Col1a1, Dcn, TNC, Scx mRNA and protein were detected by qPCR (A) and WB (B). Note: KIT-NC:KIT no-load group; Lvx-KIT: KIT overexpression group; si-KIT-NC: KIT interference no-load group;si-KIT: KIT interference group. “*” means p<0.05, “**” means p<0.01, “***” means p<0.001, the significant difference in the figure Sex was compared to the KIT-NC group.

**Fig 14.**
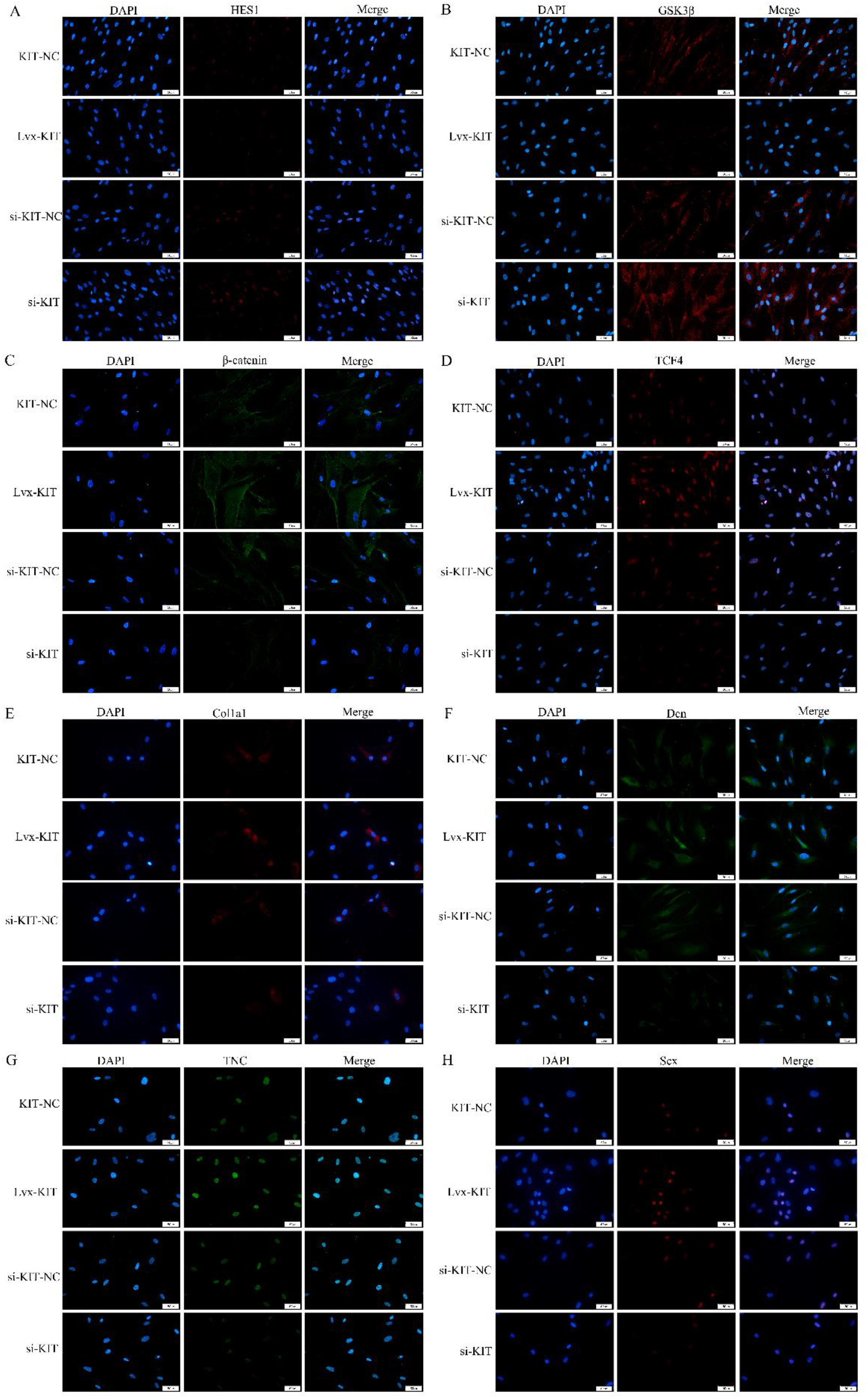
Immunofluorescence detection of protein expression levels of HES1, GSK3β, β-catenin, TCF4, Col1a1, Dcn, TNC and Scx. Note: KIT-NC:KIT no-load group; Lvx-KIT: KIT overexpression group; si-KIT-NC: KIT interference no-load group;si-KIT: KIT interference group. Scale: 50 μm.

**Fig 15.**
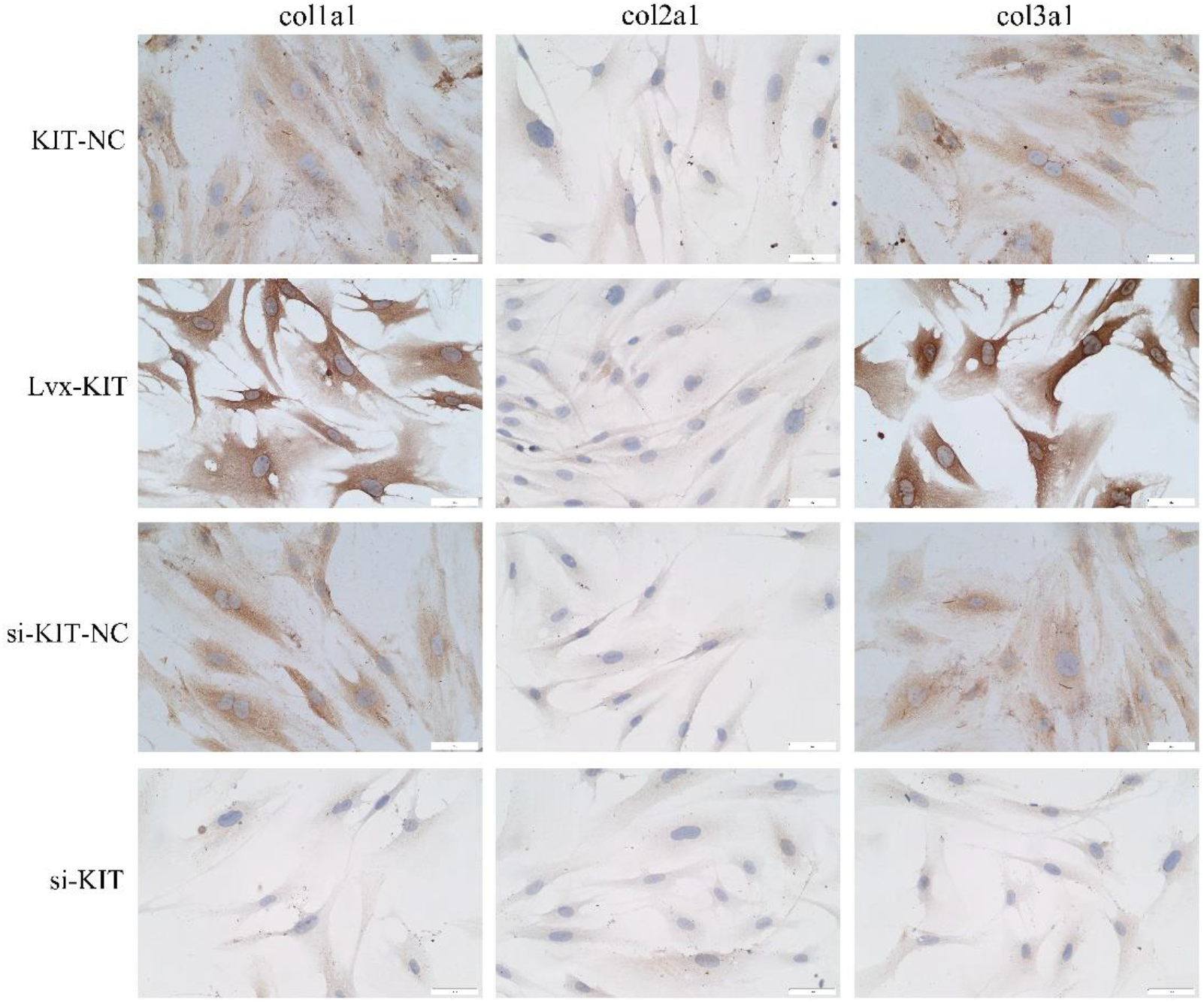
Expression levels of collagen Col1a1, Col2a1 and Col3a1 were detected by immunohistochemistry. Note: KIT-NC:KIT no-load group; Lvx-KIT: KIT overexpression group; si-KIT-NC: KIT interference no-load group;si-KIT: KIT interference group. Scale: 50 μm.

### Overexpression of KIT gene in transplanted rats TDSC promotes implant ligamentalization and tendon-bone healing

Beginning on the 14th day after modeling, the Lequesne MG behavioral scores of each group had a downward trend, especially the gene overexpression group had the most obvious downward trend after treatment. After 42 days of treatment, the score was significantly lower than the overexpression null group (Fig.16A). Samples were collected at 6-time points on the 3rd, 7th, 14th, 21st, 28th, and 42nd days of surgery. With the extension of time, the ligaments and joints of the reconstructed and repaired grafts are more and more closely integrated, and occasionally the joints of rats are infected (Fig.16C).

**Fig 16.**
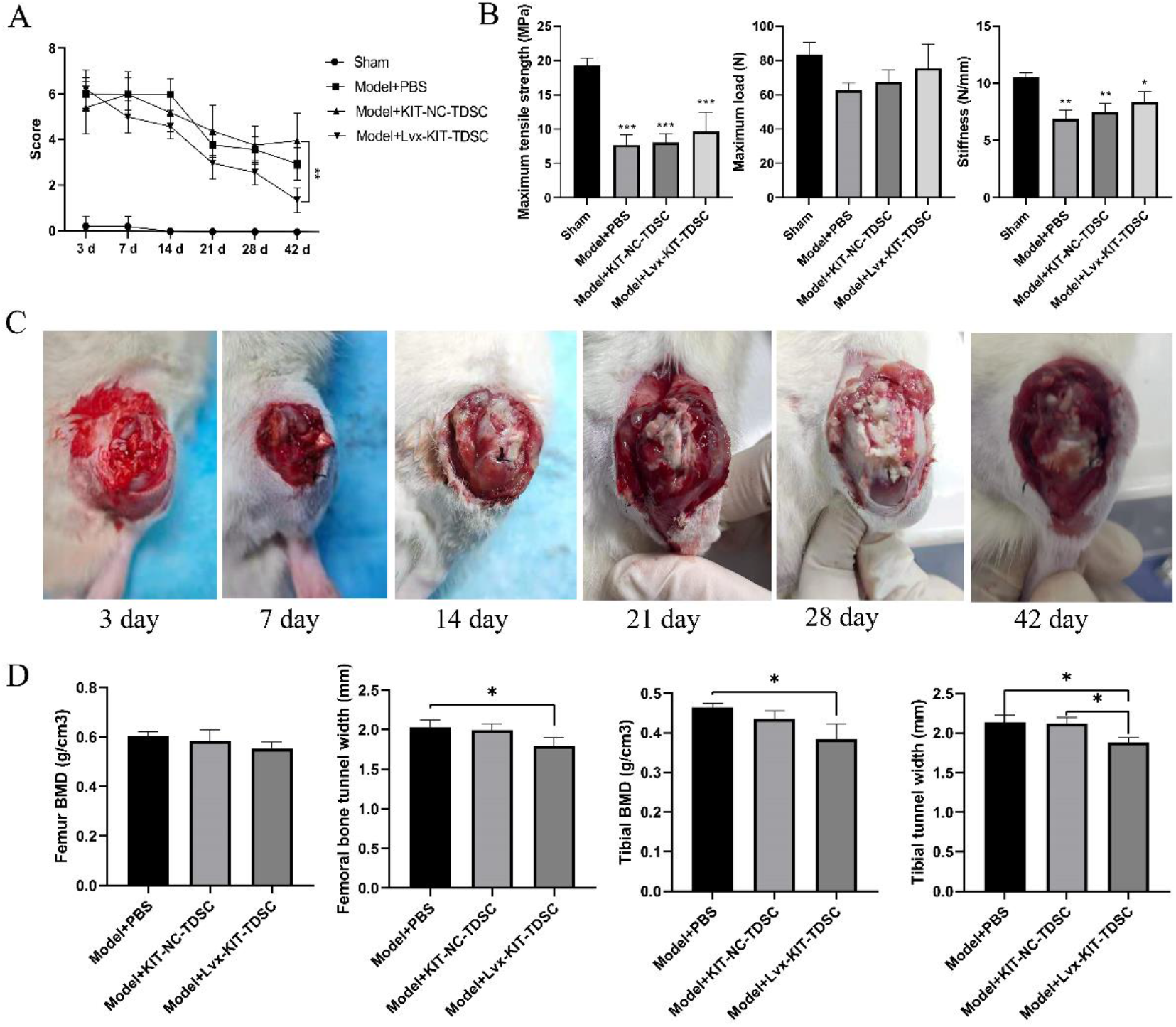
Treatment of implant ligamentization and tendon-bone healing by overexpressing TDSC in transplanted rat KIT gene to induce tendon differentiation in vivo. (A) Lequesne MG behavior scores; (B) The mechanical properties of knee joint were tested after 42 days of operation; (C) The materials were taken at 3, 7, 14, 21, 28 and 42 days after operation to observe the combination of ligament and joint; (d) Micro CT detection. Note: “*” means p<0.05, “**” means p<0.01, “***” means p<0.001.

The mechanical properties of the knee joint on the 42nd day of operation were tested. Compared with the sham operation group, the maximum tensile strength (MPa), maximum load (N) and stiffness (N/mm) of the other groups were significantly reduced (Fig.16B). However, the KIT overexpression TDSC group increased compared with the model group and the overexpression control group (p>0.05). Micro CT detection showed that the width of the bone marrow tract in the overexpressed TDSC group was significantly narrower than that in the model group after repair, and the cancellous bone density in the repaired area was significantly lower than that in the model group (Fig.16D).

HE staining showed that in the sham-operated group in 6 time periods, the fibers of the tendon in the anterior fork tissue were arranged in a tight and orderly manner, and no hemorrhage and inflammatory cells were found. On the 3rd day of operation, the fibers in the model group were loosely arranged and disordered, filled with inflammatory cells, and a small amount of dissolved muscle cells were seen; in the overexpressed no-load TDSCs, the fibers were loosely arranged and infiltrated by a large number of inflammatory cells; in the KIT gene-overexpressed TDSCs, the fiber arrangement was looser than that of the normal group, the bleeding was severe, and the infiltration of inflammatory cells was visible. Compared with the 3rd day, on the 7th day of operation, the fibers in the model group were loosely arranged and disordered, and a large number of inflammatory cells were infiltrated, but the number was reduced compared with that on the 3rd day, and bleeding was seen; Inflammatory infiltration and deposition of adipocytes were observed at the same time; in the TDSC group with KIT gene overexpression, bleeding was rare, inflammatory cell secretion serous fluid was seen at the edge, and some inflammatory cells infiltrated in the fibrous tissue.On the 28th and 42nd day of surgery, the fibers in the model group were loosely arranged, with a small amount of inflammatory cell infiltration, neovascularization, and a small amount of fibroblasts. There were many new blood vessels and a small number of fibroblasts. In the TDSC group with KIT gene overexpression, the fibers were arranged in a more compact and orderly manner, with more fibroblasts and occasional inflammatory cells. With the prolongation of treatment time, compared with other groups, the recovery effect of KIT gene overexpression TDSC group is the best. (Fig. 17)

**Fig 17.**
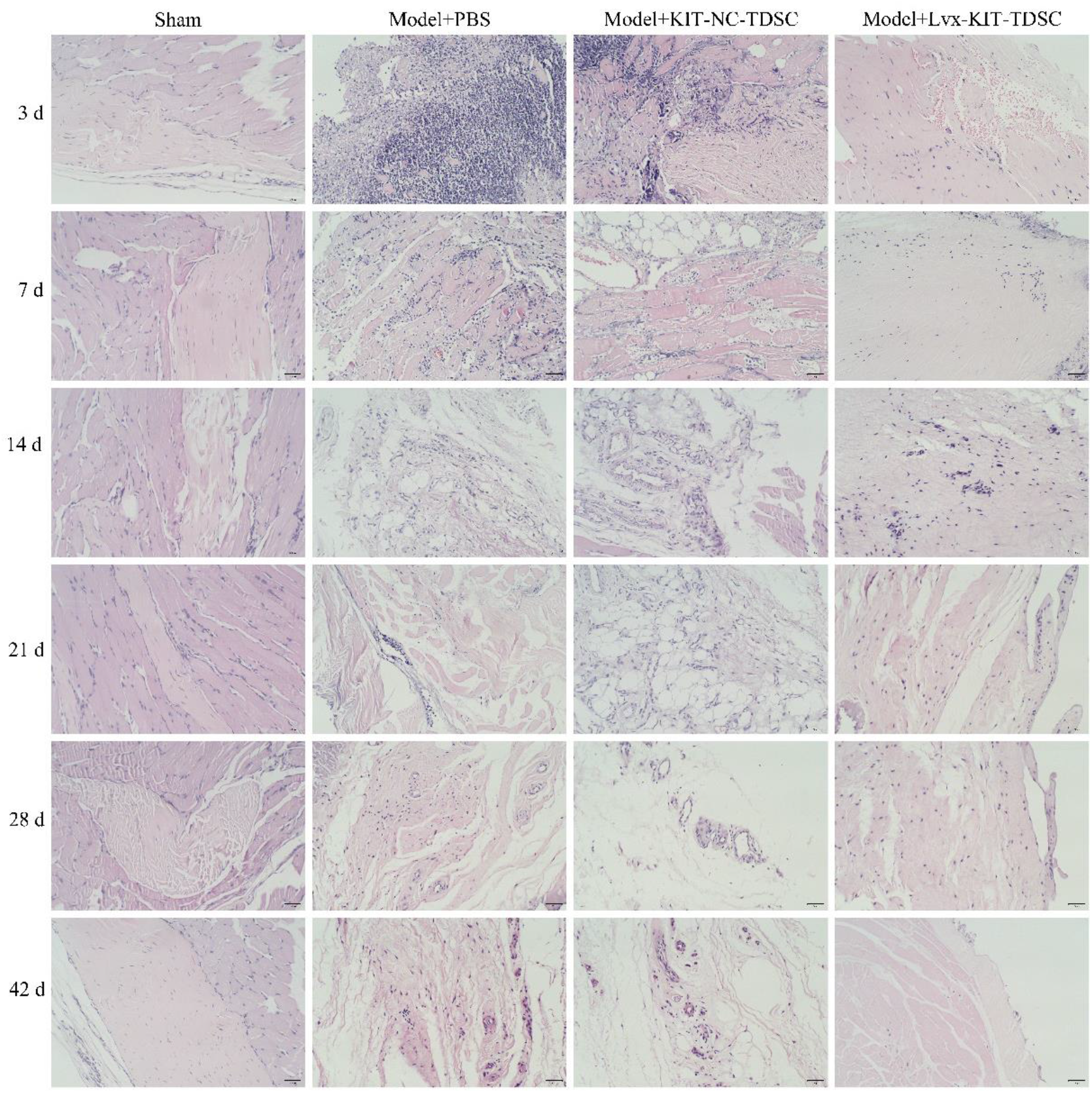
HE staining observed the recovery of the sham-operated group, model group, model group + no-load TDSC group and model group + KIT overexpression TDSC group after 3 days, 7 days, 14 days, 21 days, 28 days and 42 days of treatment Condition. Scale: 100 μm.

qPCR and WB detection showed that the mRNA and protein expressions of tendon markers and transcription factors Col1a1, Dcn, TNC, and Scx were significantly increased in the KIT overexpression TDSC group; The mRNA and protein expressions of the expected pathway proteins HES1, GSK3β, β-catenin and TCF4 were consistent with the expected results (Fig.18A,B). The results of immunofluorescence detection were consistent with the results of qPCR and WB detection (Fig.19).

**Fig 18.**
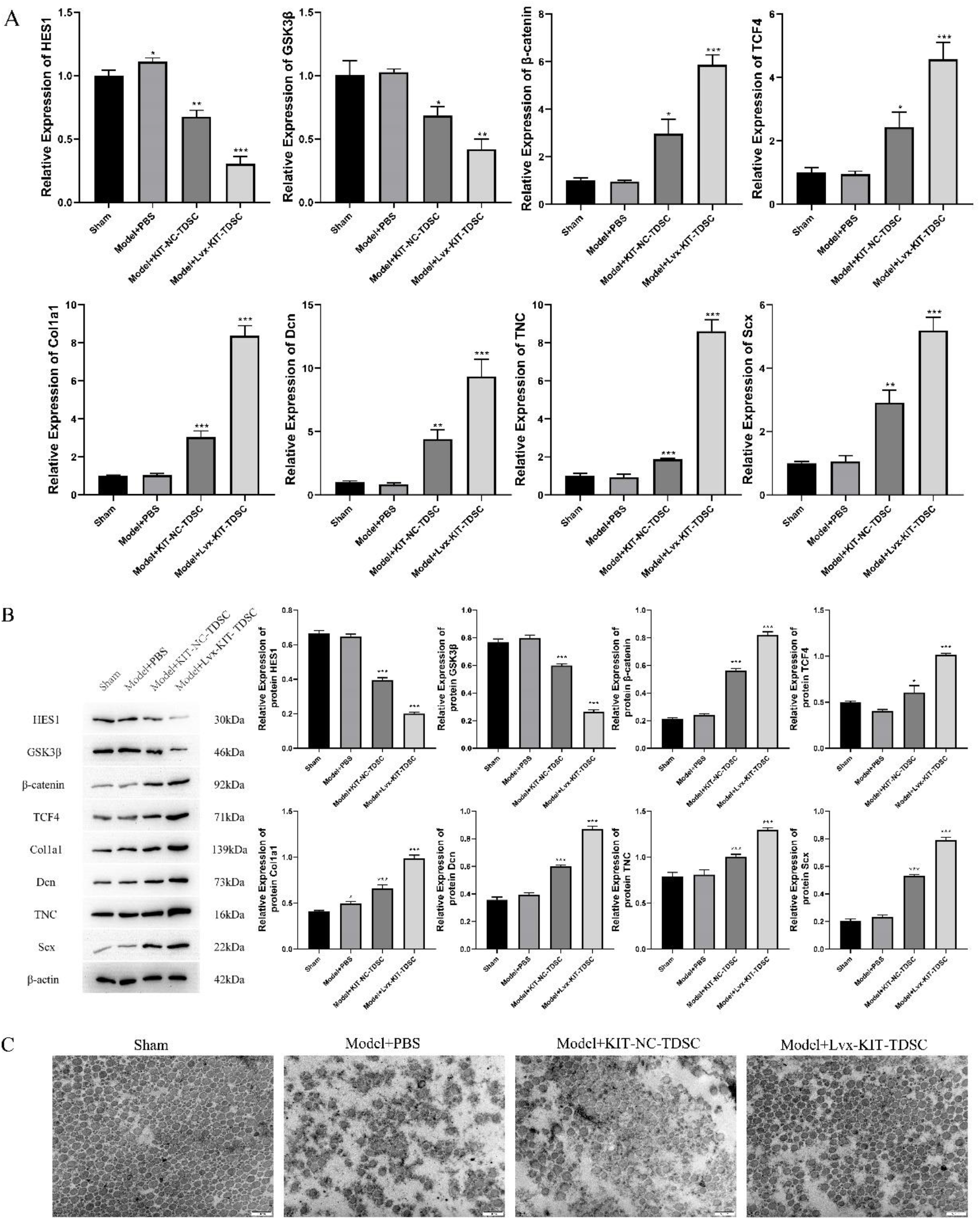
(A) qPCR and (B) WB were used to detect mRNA and protein expression of HES1, GSK3β, β-Catenin, TCF4, COL1A1, DCN, TNC and SCX; (C) Transmission electron microscope observation. Note: “*” means p<0.05, “**” means p<0.01, and “***” means p<0.001, the significant difference in the figure Sex was compared to the Sham group. Scale: 500 nm.

**Fig 19.**
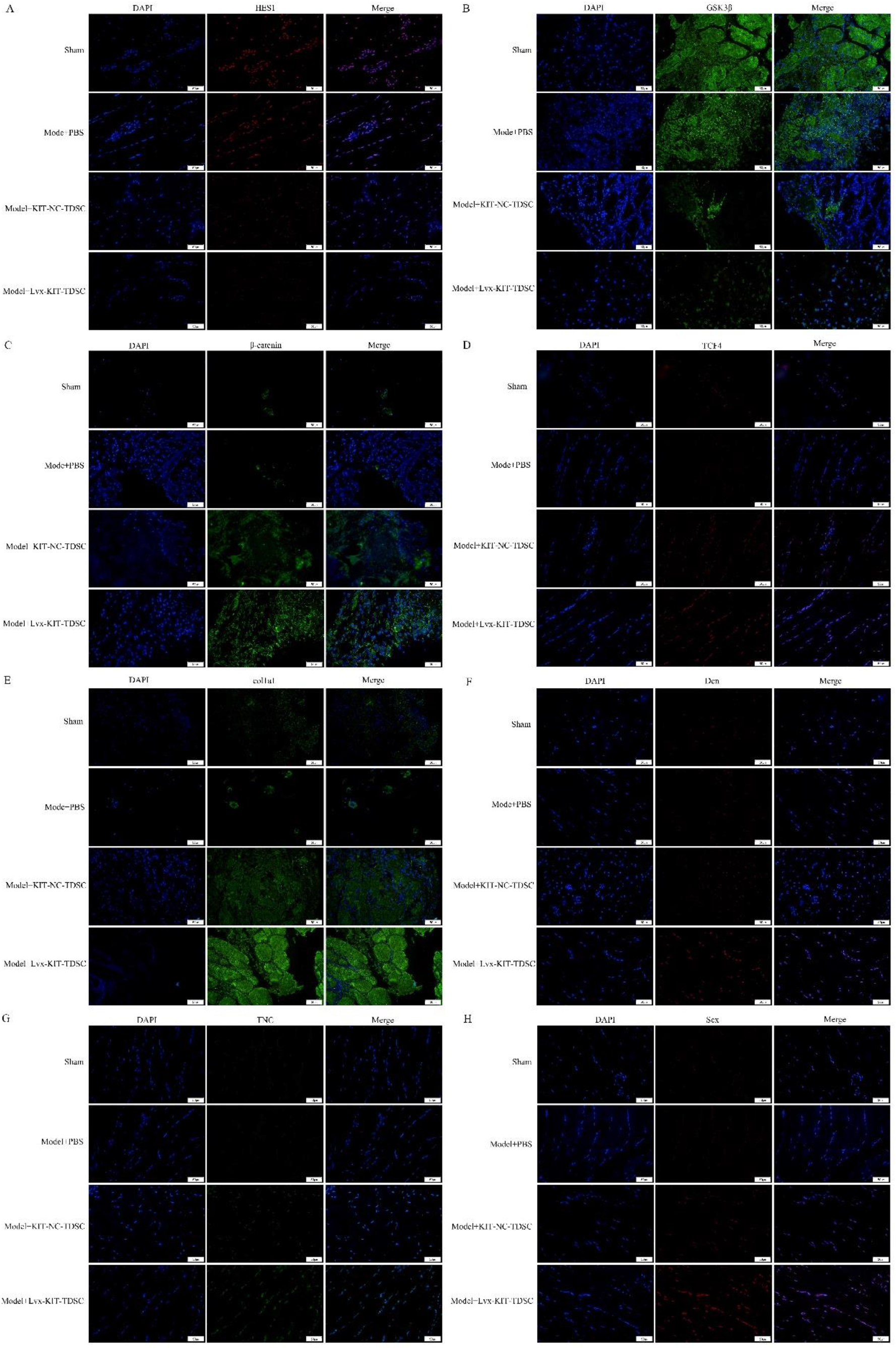
Immunofluorescence detection of protein expression levels of HES1, GSK3β, β-catenin, TCF4, Col1a1, Dcn, TNC, and Scx. Scale: 50 μm.

The results of transmission electron microscope showed that compared with the normal group, the fibers in the model group were loosely arranged and the number of fibers was less, and the number of large-diameter fibers and small-diameter fibers in the model group were lower than those in the sham operation group. The situation of the unloaded TDSC group was similar to that of the model group, and the number and compactness of fibers were both poor. In the group treated with KIT gene overexpression TDSC, the number of fibers increased significantly, the arrangement of fibers was closer, and the number of large-diameter fibers and small-diameter fibers increased significantly (Fig.18C).

Repair and remodeling of CTGF involve not only the reproduction and activity of chondrocytes but also the formation of collagen fibers and abrasive substances. Both Col1a1 and Col2a1 are important natural resources for tissue regeneration and wound healing [22]. Studies have found that CTGF stimulates the production of Col1a1 and Col2a1 in the injured ACL, which are required for ACL reconstruction and ligament recovery [23]. Immunohistochemical detection showed that with the increase in treatment days, the expression of collagen proteins Col1a1, Col2a1 and Col3a1 in the KIT overexpression TDSC group increased significantly (Fig.20).

**Fig 20.**
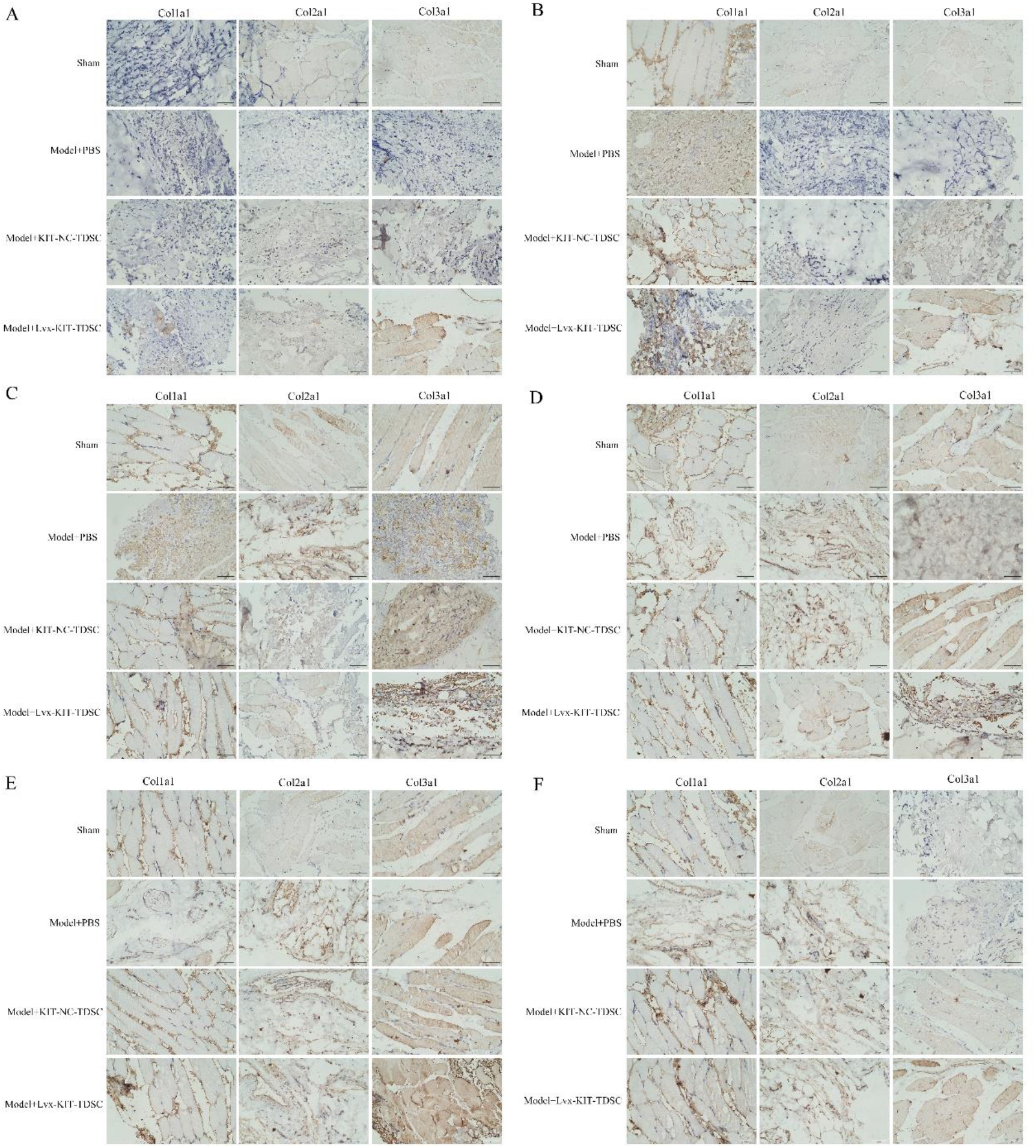
Expression levels of collagen Col1a1, Col2a1 and Col3a1 were detected by immunohistochemistry. Scale: 50 μm.

## Discussion

At present, ACL reconstruction is the most effective way to treat ACL injury. The quality of tendon-bone healing after the operation is the key to the success of the operation. If the healing speed of the tendon-bone interface is slower than that of the bone interface, the operation will fail [24]. Stem cell therapy is a research hotspot in the treatment of tendon injury. BMSC, adipose-derived stem cell (ADSC), and embryonic stem cell (ESC) are all potential treatment methods for tendon injury. Compared with other stem cells, TDSCs may be a promising source of therapeutic cells for better and earlier musculoskeletal repair, including tendon repair [14].

TDSC, a new type of pluripotent stem cell-derived from tendon tissue, has the potential for self-renewal and multidirectional differentiation and has a broad prospect of clinical application [25]. The primary isolated hTDSC have uniform morphology and stable growth. The cell surface markers CD44, CD90, and CD105 are positive and can differentiate into three lines under corresponding induction conditions. They have typical stem cell characteristics.

The study of tendon-bone interface healing in rabbits after ACL reconstruction stimulated by CTGF found that CTGF could promote the proliferation and differentiation of bone progenitor cells, adhesion and migration of osteoblasts, vascular differentiation, and formation of myofibroblasts [26,27]. The mechanism of action of CTGF involves important interactions with other regulatory signals such as Wnt, BMP, and IGF, acting through binding peptides or their receptors [28–31]. During TDSC tendon differentiation induced by CTGF, the protein expression in cells changed significantly. Go and KEGG analysis showed that there were some undetected or unexplained signal pathways related to tendon differentiation in cells, which proved that CTGF may promote TDSC tendon differentiation through some signal regulation pathways, to promote the recovery of the ACL.

The analysis of immunoprecipitation and mass spectrometry data showed that there was a direct interaction between kit and CTGF. KIT uses stem cell growth factor (SCF) as its ligand, so it is called stem cell growth factor receptor (SCFR), which plays an important role in stem cell maintenance and differentiation [32, 33]. The activation of the SCF / c-kit signal transduction pathway is usually related to cell proliferation, migration and survival [34]. The human TDSC tendon induction model in vitro confirmed that the expression of KIT gene was positively correlated with the expression of tendon markers COL1A1, DCN, TNC and SCX and the differentiation of stem cells.

The changing trend of HES1, GSK3β, β-Catenin and TCF4 with the change of KIT gene expression is consistent with the expected change. HES1 is a transcription factor that affects cell proliferation and differentiation during embryogenesis [35]. Studies have shown that HES1 affects the maintenance of certain stem cells and ancestral cells [36, 37]. The results of Chip-qPCR and double luciferase reporter gene detection proved that GSK3β is a downstream target gene regulated by HES1. Glycogen synthase kinase-3β (GSK-3β)is an evolutionarily conserved serine/threonine kinase that plays a role in many cellular processes, including cell proliferation, DNA repair, cell cycle, signal transduction and metabolic pathways. GSK-3β participates in a variety of signaling pathways, including Wnt/ β-Catenin, PI3K / PTEN / Akt and notch [38]. It is proved that CTGF mediates Wnt / β-Catenin signaling pathway by directly interacting with KIT to regulate transcription factor HES1 promotes the tendon differentiation of tendon stem cells.

KIT gene overexpression and KIT gene interference induced tendon differentiation of rat TDSC stable transfer under CTGF to verify the efficacy of implant ligamentization and tendon-bone healing. The experimental results show that after repair surgery, TDSC with KIT gene overexpression is used for treatment. Through the detection of animal behavior scores, mechanical properties and HE staining tissue observation, it is found that this method has a better recovery effect than other methods. The detection of key proteins of related signal pathways and tendon markers in each group also obtained a trend similar to the experimental results of molecular mechanisms at the cell level, which verified the previous experimental results at the animal level. In this study, it was confirmed in hTDSC and animal experiments that the modified hTDSC can promote graft ligamentation and tendon-bone healing after ACL reconstruction.

## conclusion

The formal separation of hTDSC has uniform morphology and stable growth. The cell surface markers CD44, CD90 and CD105 are positive, and can differentiate into three lines under corresponding induction conditions. They have typical stem cell characteristics. CGTF membrane receptor KIT can be adjusted, which can combine more CTGFs to inhibit the downstream HES1. After HES1 is suppressed, the transcription of GSK3β is inhibited, the expression volume decreases, and the corresponding β-catenin inhibited by GSK3β is activated, thereby activating the downstream transcription factor TCF4 and promoting the tenderization of tendon stem cells.

